# ENKD1 is a centrosomal and ciliary microtubule-associated protein important for primary cilium assembly and Hedgehog signaling

**DOI:** 10.1101/2021.07.20.453048

**Authors:** Fatmanur Tiryaki, Jovana Deretic, Elif Nur Firat-Karalar

**Affiliations:** Department of Molecular Biology and Genetics, Koç University, Istanbul, Turkey 34450; Koç University School of Medicine, Istanbul, Turkey 34450

**Keywords:** ENKD1, microtubules, centrosome, primary cilium

## Abstract

Centrioles and cilia are conserved, microtubule-based structures critical for cell function and development. Their structural and functional defects cause cancer and developmental disorders. How microtubules are organized into ordered structures by microtubule-associated proteins (MAPs) and tubulin modifications is best understood during mitosis but is largely unexplored for the centrioles and the ciliary axoneme, which are composed of remarkably stable microtubules that maintain their length at steady state. In particular, we know little about the identity of the centriolar and ciliary MAPs and how they work together during the assembly and maintenance of the cilium and centriole. Here, we identified Enkurin domain containing 1 (ENKD1) as a component of the centriole wall and the axoneme in mammalian cells, and showed that it has extensive proximity interactions with these compartments and MAPs. Using *in vitro* and cellular assays, we found that ENKD1 is a new MAP that promotes microtubule polymerization and regulates microtubule organization and stability. Consistently, overexpression of ENKD1 increased tubulin polymerization and acetylation and disrupted microtubule organization. Cells depleted for ENKD1 were defective in ciliary length and content regulation and failed to respond to Hedgehog pathway activation. Together, our results establish ENKD1 as a new centriolar and ciliary MAP that regulate primary cilium structure and function, and advances our understanding of the functional and regulatory relationship between MAPs and the primary cilium.

## Introduction

Microtubules are dynamic cytoskeletal filaments assembled by polymerization of alpha- and beta-tubulin heterodimers into hollow cylinders. Although they are evolutionarily highly conserved, microtubules assemble functionally and structurally diverse cellular structures such as the bipolar mitotic spindle, centrioles, cilia and flagella. Given the critical roles of these structures for proper cellular function and development, defects in their biogenesis and function cause various human diseases ranging from cancer to developmental disorders (Godinho and Pellman, 2014; Higgins et al., 2019; O’Neill et al., 2018; Reiter and Leroux, 2017). The major microtubule-organizing center (MTOC) in most animal cells is the centrosome, which is composed of two centrioles and associated pericentriolar material (PCM) (Akhmanova and Steinmetz, 2015; Paz and Luders, 2018; Sanchez and Feldman, 2017; Wu and Akhmanova, 2017). Through concentrating gamma-tubulin, tubulin dimers and microtubule-associated proteins (MAPs), PCM supports microtubule nucleation, polymerization and organization (Luders, 2012; Woodruff et al., 2014). In interphase cells, centrosomes form radial microtubule arrays that plays important roles in determining cell shape and providing tracks for molecular motor-based intracellular transport. In dividing cells, centrosomes contribute to the assembly of the bipolar spindle that is required for equal segregation of chromosomes and positioning the cell division plane. In response to quiescence and differentiation, one of centrioles of the centrosome nucleates the formation of cilia (Mirvis et al., 2018; Sanchez and Dynlacht, 2016). In contrast to the cytoplasmic microtubules that undergo dynamic instability, centriolar and ciliary microtubules are highly stable and they maintain their length at steady state. Our understanding of the shared and distinct mechanisms by which distinct microtubule-based structures are assembled, maintained and dynamically altered is incomplete.

Centrioles are evolutionarily conserved microtubule-based cylindrical structures built from nine radially arranged compound microtubules that are polarized along their long axis (Wang and Stearns, 2017). During ciliogenesis, centrioles act as basal bodies to template the nine axonemal microtubule doublets of the primary cilium and motile cilia (Mirvis et al., 2018; Pedersen et al., 2012). The axoneme serves as the structural backbone of the cilia and the track for bidirectional movement of ciliary cargoes such as signaling molecules and structural components. Most cells in human body bear a surface-exposed primary cilium, which compartmentalizes signaling molecules and acts as a sensory hub for a wide range of signaling pathways required for embryonic development and tissue homeostasis (Nachury, 2014; Nachury, 2018; Wheway et al., 2018). For example, Hedgehog signaling components Patched (PTCH1) receptor, Smoothened (SMO), GPR161, SUFU and Gli dynamically traffic into and out of the primary cilium in response to Hedgehog (Hh) ligands and turn on Hh target gene transcription (Bangs and Anderson, 2017). Although sensory functions of the primary cilium require tight regulation of the structure, length and stability of the centriolar and axonemal microtubules, how this is achieved in cells remains poorly understood.

The diversity of microtubule-based structures and their dynamic adaptations in response to different stimuli have been shown to be regulated by multiple mechanisms. First, microtubules interact with a diverse array of microtubule-associated proteins (MAPs) that regulate microtubule dynamics, organization and stability (Bodakuntla et al., 2019; Conkar and Firat-Karalar, 2020; Goodson and Jonasson, 2018). One group of MAPs bind to the microtubule lattice and mediate its interactions with other proteins or stabilize it. Another group binds to microtubule ends and regulate dynamic instability and their attachment to other cellular structures (Akhmanova and Steinmetz, 2015). Finally, molecular motors are responsible for directed motility of cargoes along microtubules or for force generation. In addition to MAPs, the incorporation of different tubulin isoforms to the microtubules and their posttranslational modifications (i.e. acetylation, polyglutamylation) contribute to the heterogeneity of microtubules (Chakraborti et al., 2016; Gadadhar et al., 2017; Janke and Magiera, 2020). Extensive progress has been made in elucidating how MAPs and tubulin code regulate interphase and mitotic microtubules. However, we know little about the identity of the centriolar and ciliary MAPs and their interactions with the axoneme and with each other, and their roles in cilia organization and function.

The stability and dynamic behavior of centriolar and axonemal microtubule are different from that of cytoplasmic microtubules, suggesting that they are differentially regulated. Both centriolar and axonemal microtubules are remarkably stable and maintain their length when they reach steady state (Conkar and Firat-Karalar, 2020; Wang and Stearns, 2017). In contrast to centrioles, axonemal microtubules undergo constant polymerization and depolymerization even at steady state. High-resolution molecular structure of the primary cilium has been described by recent studies that exploited expansion microscopy and cryo-electron tomography (Conkar and Firat-Karalar, 2020; Katoh et al., 2020; Kiesel et al., 2020; Le Guennec et al., 2020; Sun et al., 2019). Although these studies revealed major differences between the three-dimensional organization of axonemal microtubules in primary and motile cilia, the molecular basis of these differences is not understood. The number and type of MAPs that localize the axonemes are likely to contribute to these differences. Consistently, new axonemal protein densities have been defined by structural studies, but the molecular make up of these densities and how they contribute to axoneme structure and function remains unknown (Kiesel et al., 2020; Sun et al., 2019). To address these questions, it is essential to first complete the parts list for the ciliary MAPs and then to elucidate how MAPs and microtubules cooperate to assemble the ciliary axoneme at the right size and architecture.

The proteomes of the centrioles and cilia have been largely identified using a combination of different approaches such as proteomics, transcriptomics and genomics (Breslow and Holland, 2019; van Dam et al., 2019). To define and characterize new MAPs required for the structure and function of centrioles and cilia, we examined published datasets and identified Enkurin domain containing 1 (ENKD1/C16orf48) as a putative centrosomal and/or ciliary MAP based on several lines of evidence. ENKD1 was originally identified by a high-throughput imaging screen for identification of new MAPs, and HA-FLAG-ENKD1 was shown to localize to interphase microtubules in this study (Fong et al., 2013). Subsequently, systematic subcellular localization analysis of 506 human proteins in human cell lines showed that endogenous ENKD1 localizes to the centrosome, whereas ENKD1-GFP localizes both to the centrosome and microtubules (Stadler et al., 2013). In addition to studies in mammalian cell lines, ENKD1 was shown to be transcriptionally upregulated during centriole duplication and motile cilia assembly in *in vitro* mouse tracheal epithelial cultures (MTECs) (Hoh et al., 2012) and was also was identified as a conserved ciliary candidate protein through evolutionary proteomics of motile cilia from sea urchins, sea anemones and choanoflagellates (Sigg et al., 2017). Finally, ENKD1 was identified as a component of the primary cilium proteome generated in kidney epithelial IMCD3 cells using cilia-APEX2-proximity labeling combined with quantitative mass spectrometry (May et al., 2021) and was described as a potential cancer biomarker (Song et al., 2021). Based on these results, we hypothesized that ENKD1 is a ubiquitous centrosomal and/or ciliary MAP. However, its precise cellular functions and mechanisms are not known.

Here, we showed that ENKD1 is a new MAP that regulates microtubule polymerization, organization and stability *in vitro* and in cells. ENKD1 localizes to the centrosome, primary cilium, and microtubules through distinct N- and C-terminal domains and has extensive proximity interactions with these structures. Ultrastucture expansion microscopy revealed localization of ENKD1 to the centriole wall, pericentriolar material and ciliary axoneme. Overexpression of ENKD1 promotes microtubule polymerization and stability, and interferes with microtubule organization. Its depletion results in shorter cilia, deregulated ciliary content and defective Hedgehog pathway activation. Collectively, our results identify ENKD1 as a MAP that regulates primary cilium biogenesis and function via its microtubule-associated activities.

## Results

### ENKD1 localizes to the centrosomes and primary cilia

ENKD1 is conserved in diverse organisms containing centrioles and cilia ranging from humans to *Trypanosomes* (Sigg et al., 2017) (Fig. S1). mRNA and protein expression profiling studies showed that ENKD1 is ubiquitously expressed across various tissues (Thul et al., 2017; Uhlen et al., 2015). Unlike ENKD1, the expression of its mammalian ortholog ENKUR was restricted to the tissues with motile cilia such as trachea and testis. In agreement with its localization, ENKUR was shown to regulate motile cilia assembly and function (Sigg et al., 2017; Sutton et al., 2004). The wide expression profile of ENKD1 suggests its functions across different cell types such as cells that assemble primary cilia or motile cilia.

To investigate cellular functions of ENKD1, we first examined the localization of endogenous ENKD1 and its fluorescent fusion proteins in mouse inner medulla collecting duct IMCD3 and human retinal pigmented epithelial RPE1 cells, which form primary cilium upon serum starvation. Polyclonal antibodies against ENKD1 revealed that it localizes to the centrosome and the primary cilia in RPE1 cells (Fig. 1A, S2A). The two polyclonal antibodies raised against different ENKD1 antigens recognized ENKD1 fusion proteins (Fig. S1, S2B), but did not work for immunofluorescence in IMCD3 cells. To assess its localization and dynamic behavior, we next used lentiviral transduction combined with serial dilutions and generated RPE1 and IMCD3 clonal lines stably expressing express mNeonGreen (mNG)-ENKD1 at low and high expression levels, as determined by immunoblotting using antibodies against ENKD1 and mNG for RPE1 cells (Fig. S2C). In low-expressing RPE1 and IMCD3 cells, mNG-ENKD1 localized to the centrosome during interphase, to the spindle poles during mitosis and to the centrosomes and central spindle during cytokinesis (Fig. 1B, 1C, S2D). In high-expressing RPE1 cells, mNG-ENKD1 additionally localized to the radial microtubule array during interphase and to the bipolar spindle during mitosis (Fig. S2E, S2F). Similar centrosome and microtubule localization was observed when 6xMyc-ENKD1 was transiently expressed in RPE1 cells (Fig. 2B). Of note, ENKD1 antibody did not recognize the interphase and mitotic microtubules visualized in high-expressing stable cells, suggesting that ENKD1 localization to these structures might be transient or a consequence of ectopic expression or tagging the protein (Fig. 1A, S2A).

**Figure 1.**
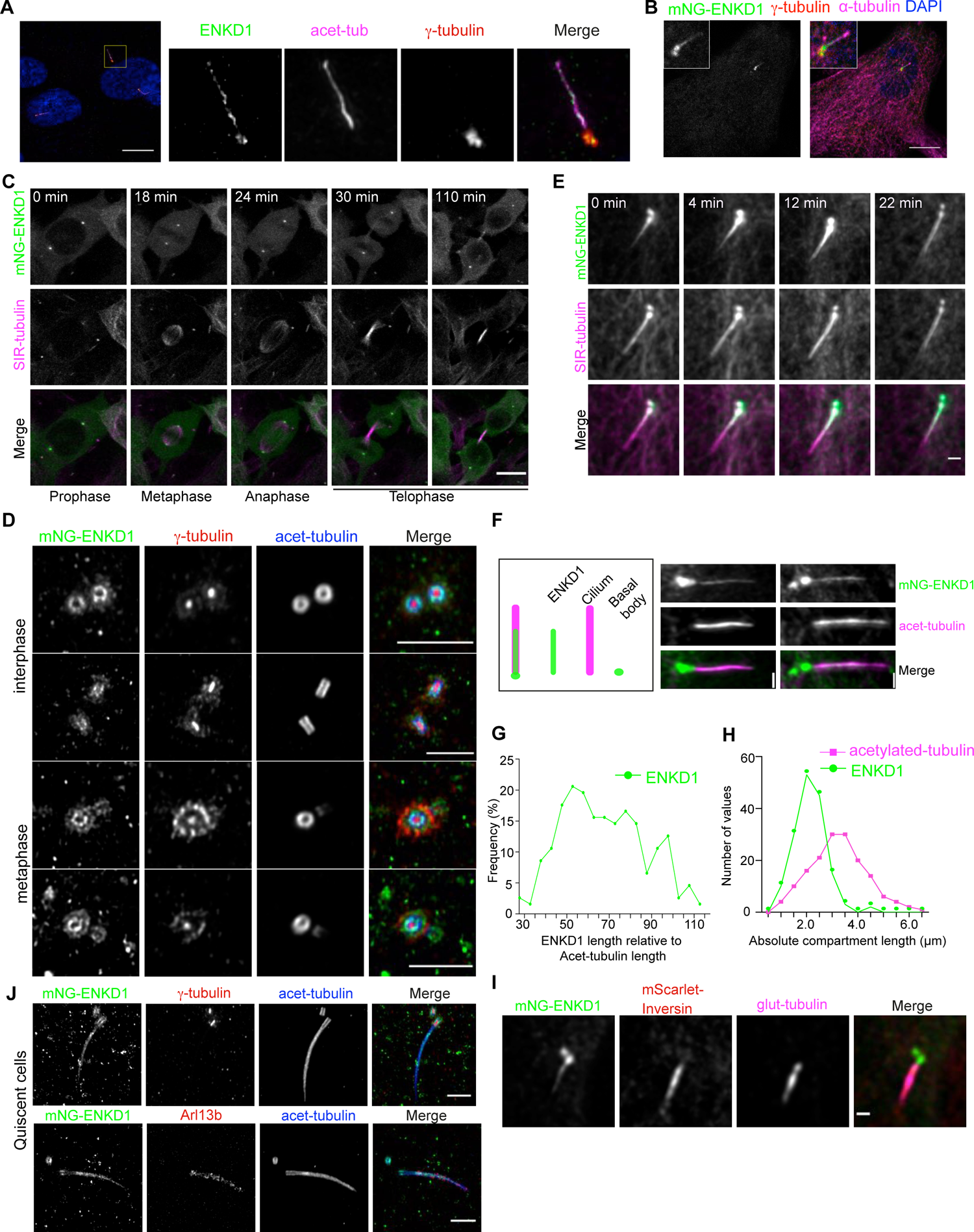
ENKD1 localizes to the centrosomes and primary cilia. (A) Endogenous localization of ENKD1 in ciliated cells. RPE1 cells were serum serum starved for 24h, fixed and stained with rat polyclonal antibody raised against full length ENKD1, `γ`-tubulin, and acetylated tubulin (acet-tub). Scale bar: 10um (B) ENKD1 localization in ciliated low-expressing RPE1 mNeonGreen-ENKD1 (RPE1::mNG-ENKD1) stable cells. Cells were serum staved for 24 h, fixed with PFA and stained for gamma-tubulin (`γ`-tubulin), alpha-tubulin (`α`-Tubulin) and DAPI. Scale bar: 10 `μ`m. (C) Dynamic localization of ENKD1 throughout the cell cycle. RPE1::mNG-ENKD1 low expressing cells were stained with SiR-tubulin and analysed by time GFPse confocal imaging. Still frames from Movie 1 were shown at the indicated time points to represent ENKD1 localization to the spindle poles and central spindle. Scale bar: 10 `μ`m (D) Nanoscale localization of ENKD1 at different stages of the cell cycle. RPE1::mNG-ENKD1 low-expressing cells were expanded with U-ExM, and stained with antibodies against ENKD1, gamma-tubulin to mark the pericentriolar material, and acetylated-tubulin to mark the centriole and cilium microtubules. Top two panels represent interphase localization; bottom two panels represent metaphase localization. Scale bar: 1 `μ`m (E) Constitutive localization of of ENKD1 at the primary cilium. Low-expressing RPE1::mNG-ENKD1 cells were serum staved for 24 h, stained with SiR-tubulin and imaged for 22 min under serum starvation conditions. Still frames from Movie 2 were shown at the indicated time points to represent localization of ENKD1 to the primary cilium during the time course. Scale bar: 1 `μ`m (F) Analysis of ENKD1 distribution along the primary cilium. Schematic illustrates the color code and the method for quantification of ENKD1 distribution along individual primary ciliu. RPE1::mNG-ENKD1 cells were serum starved for 24 h, fixed with PFA and stained for acetylated-tubulin (acet-tub). mNG-ENKD1 (green) signal either spans the whole cilium (left panel) or proximal part of the cilium (right panel). Scale bar: 1 `μ`m (G) Distribution of relative ENKD1 length. Length of ENKD1 signal was normalized relative to cilium length: [length ENKD1]/ [length AcTub]. Only ENKD1 positive cilia were included for the analysis. N=182 (H) Distribution of absolute ENKD1(green) and acetylated tubulin(magenta) lengths measured in `μ`m by confocal imaging of RPE1::mNG-ENKD1 cells stained with anti-acetylated-tubulin antibody. Only ENKD1 positive cilia are included. Mean ENKD1 length is 2.1 `μ`m (SD =0.60, n= 158) and mean acetylated-tubulin length is 3.3 `μ`m (SD = 1.1, n= 158). (I) ENKD1 localization at the primary cilium relative to the inversin compartment. RPE1::mNG-ENKD1 cells stably expressing mScarlet-Inversin were serum starved for 24 h, fixed with PFA and stained for polyglutamylated-tubulin (glut-tub). Scale bar: 1 `μ`m (J) High-resolution ciliary localization of ENKD1 in quiescent ciliated cells. RPE1::mNG-ENKD1 cells were serum starved for 24h and expanded with U-ExM, and stained for ENKD1, gamma-tubulin to mark the pericentriolar material, and acetylated-tubulin to mark the centriole and cilium microtubules. Scale bar: 1 `μ`m

**Figure 2.**
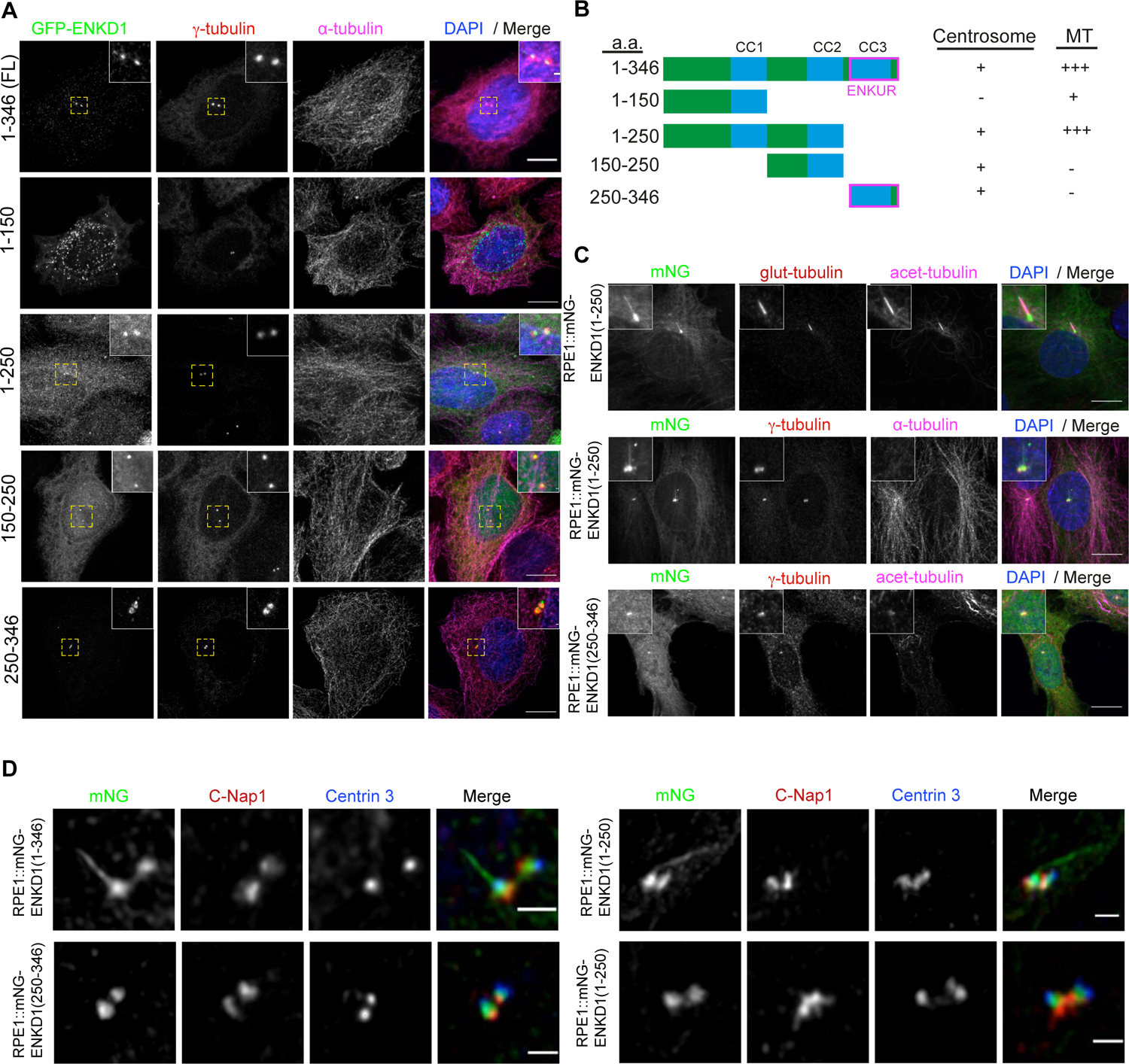
Localization analysis of ENKD1 deletion mutants. (A) Localization of GFP-fusions of ENKD1 and its deletion mutants in transiently-transfected low-expressing RPE1 cells. Hela cells were with the indicated constructs, fixed and stained for GFP, `γ`-tubulin and microtubules (α-tubulin). Centrosome localization was observed for ENKD1 FL (1-346), (1-250), (150-250) and (250-346) fragments. Scale bar: 10 `μ`m (B) Schematic representation of ENKD1 full length (FL) and deletion mutants and summary of their localization to the centrosome, and their interaction or association with microtubules. Centrosome localization and microtubule association were determined by analysing Hela cells transiently transfected with GFP-ENKD1 constructs and staining fixed cells for GFP, `γ`-tubulin and microtubules. Numbers indicate amino acid positions, CC1, CC2, and CC3 indicate coiled coil domains and ‘ENKUR’ indicates the ENKUR domain. (C) Localization analysis of RPE1 cells stably expressing mNG fusions of N-terminal (1-250) or C-terminal (250-346) ENKD1 fragments. RPE1::mNG-ENKD1 (1-250) and RPE1::mNG-ENKD1 (250-346) cells were serum starved for 24hr, fixed with PFA and stained for polyglutamylated-tubulin and acetylated tubulin, or `γ`-tubulin and (α-tubulin; `γ`-tubulin and acetylated-tubulin. RPE1::mNG-ENKD1 (1-250) localizes to the basal body and primary cilium, RPE1::mNG-ENKD1 (250-346) localizes to the centrosome. Scale bar: 10 `μ`m (D) High resolution localization of RPE1::mNG-ENKD1, RPE1::mNG-ENKD1 (1-250), RPE1::mNG-ENKD1(250-346) cells by Hyvolution analysis. Cells were fixed and stained with proximal (C-NAP1) and distal (Centrin 3) centriole markers. All fragments localized to the middle proximal region of centrioles. FL and (1-250) fragments also localized to the primary cilium. Scale bar: 1 `μ`m

Next, we determined the subcentrosomal localization of ENKD1 in order to generate hypothesis about its function. To this end, we analyzed its localization at different stages of the cell cycle using ultrastructure expansion microscopy (U-ExM) that allows nanoscale mapping of proteins at the centrosomes and cilia (Gambarotto et al., 2019). RPE1::mNG-ENKD1 cells were stained for acetylated tubulin, which marks the centriole wall and gamma-tubulin, which marks the pericentriolar material and the centriole lumen (Schweizer et al., 2020). ENKD1 localized to resolvable rings at the outer, proximal part of the centrioles in interphase and metaphase cells and the diameter of its rings were smaller than that of gamma-tubulin rings (Fig. 1D). In addition to its localization at the PCM, ENKD1 co-localized with acetylated tubulin both in interphase and mitotic cells, which identifies it as a component of the centriole wall and suggests potential association with centriole microtubules (Fig. 1D). These results show that ENKD1 has distinct centrosomal pools at the centriole and PCM, which might be involved in different functions.

In serum-starved RPE1 / IMCD3::mNG-ENKD1 cells, ENKD1 localized to the basal body and the primary cilium (Fig. 1A, 1E, S2G). Strikingly, ENKD1 did not localize along the cilium length in all cells, instead it was restricted to the proximal end of the cilium in a fraction of cells (Fig. 1F). In RPE1 cells, the mean length of the ENKD1 ciliary region was 2.14 µm, corresponding to the %69.9 of the cilium length, which was quantified by using the axonemal acetylated tubulin as a proxy (Fig. 1F, 1G, 1H). To further define the ciliary sub-compartmentalization of ENKD1, we determined its localization relative to the proximal axoneme modified by polyglutamylation as well as the inversin compartment, which was shown to vary in length from cilium to cilium (Bennett et al., 2020; Sang et al., 2011). mNG-ENKD1 did not co-localize with polyglutamylated tubulin and mScarlet-INVS, suggesting that ciliary proteins or modifications other than polyglutamylation and inversin might regulate its ciliary incorporation (Fig. 1I). Given that ENKD1 localizes to the centriole wall, we next asked whether it also localizes to the ciliary axoneme. To this end, we performed U-ExM in ciliated RPE1::mNG-ENKD1 cells stained for the axonemal marker acetylated tubulin and the ciliary membrane marker Arl13b. ENKD1 localized along the ciliary axoneme but not at the ciliary membrane (Fig. 1J), which is reminiscent of the localization of the structural components of the cilia such as the ciliary MAPs (i.e. CCDC66, CSPP1) (Conkar et al., 2017; Conkar and Firat-Karalar, 2020; Frikstad et al., 2019).

Finally, we examined ENKD1 dynamics at the basal body and the primary cilium. First, we performed live imaging of serum-starved IMCD3/RPE1:mNG-ENKD1 stained with SIR-tubulin. After its entry into the primary cilium, mNG-ENKD1 remained there during about 30 min of imaging (Fig. 1E, S2G). Such constitutive ciliary localization is different from the ciliary dynamics of proteins implicated in signaling such as the BBSome complex and Rilp2, which transiently localize to cilia as part of their function (Nachury et al., 2007; Schaub and Stearns, 2013). Next, to determine whether ENKD1 is stably associated with basal body and cilia or is dynamic and exchangeable, we performed Fluorescence Recovery After Photobleaching (FRAP) experiments in ciliated RPE1::mNG-ENKD1 cells. About 79.68% of its basal body pool (t1/2: 21.86±0.8 sec) and 74.68% of its ciliary pool (t1/2: 31.74±1.5 sec) recovered rapidly after photobleaching (Fig. S2H). The presence of both mobile and immobile ciliary pools of ENKD1 suggest that it might function as a structural component and/or as a component of the ciliary trafficking and signaling machineries. Collectively, results from our analysis of ENKD1 localization and dynamics suggest that ENKD1 binds to microtubules and functions in centrosomal and/or ciliary processes.

### ENKD1 localizes to the centrosome, cilium and microtubules through distinct regions

Given that ENKD1 has distinct pools within the centrosome/cilium complex, we next determined the domains required for its localization to microtubules, centrosomes and cilia. To this end, we generated GFP-fusions of ENKD1-deletion mutants and characterized them for cellular localization in cells transiently or stably expressing the fusion proteins. We designed the deletion mutants of ENKD1 based on its domain organization and evolutionary conservation profile. *Homo sapiens* ENKD1 is a 346 amino acid (a.a.) protein, which contains three predicted coiled-coil (CC) domains (CC1: 101-150 a.a, CC2: 213-251 a.a., CC3: 279-335 a.a.) and an evolutionarily conserved enkurin domain (251-343 a.a.) (Fig. S1). Multiple sequence alignment of ENKD1 orthologues from diverse organisms revealed multiple conserved regions in ENKD1 including the C-terminal “enkurin” domain, which was reported to mediate interactions with TRP channels (Sutton et al., 2004).

Immunoblotting of lysates prepared from cells expressing ENKD1 deletion constructs validated their expression at the expected size (Fig. S3A). In low/moderate-expressing RPE1 cells, GFP-ENKD1 (1-250) / (150-250) / (250-346) all localized to the centrosome (Fig. 2A, 2B, S3A, S3B). GFP-ENKD1 (1-250) and (251-346) remained associated with the centrosome upon microtubule depolymerization in nocodazole-treated cells, which confirms their bona fide centrosomal localization (Fig. S3C). GFP-ENKD1 (1-150) localized to small cytoplasmic granules distributed throughout the cytoplasm (Fig. 2A, 2B, S3A, S3B). To characterize these granules, we co-stained GFP-ENKD1(1-150) with different organelle markers and found that GFP-ENKD1 (1-150) did not co-localize with mitochondria, lysosomes, endosomes, endoplasmic reticulum, Golgi and centriolar satellites (Fig. S3D). In transiently transfected IMCD3 cells, ENKD1 deletion mutants exhibited similar localization profiles to RPE1 cells (Fig. S3E).

Because ENKD1 (1-250) and ENKD1 (250-346) localized to centrosomes upon their transient expression in cells, we chose these fragments for further characterization in stable lines and functional assays. First, we generated RPE1 cells that stably express mNG fusions of N-terminal 1-250 a.a. and C-terminal 250-346 a.a. fragments of ENKD1. The expression of the fusion proteins at the expected sizes in the stable lines were confirmed by immunoblotting of RPE1 cell extracts (Fig. S3F) Upon 24 h serum starvation of these cells, we found that mNG-ENKD1(1-250) localizes to the basal body and the centrosome whereas mNG-ENKD1(250-346) localization was restricted to the centrosome (Fig. 2C). Notably, mNG-ENKD1 (1-250) localized to the primary cilium in addition to its basal body localization, which identifies this region as the ciliary-targeting domain. Finally, we determined the subcentrosomal localization of mNG-ENKD1 (1-250) and (250-346) in the RPE1 stable lines stained for the proximal centriole marker C-Nap1 and distal centriole lumen marker Centrin 3. Like mNG-ENKD1, both fragments localized to the mid-proximal region of the centrosome between C-Nap1 and Centrin 3. (Fig. 2D). Collectively, these results show ENKD1 is targeted to the centrosome through distinct regions at its N- and C-termini and to the microtubules through its N-terminal region.

### Overexpression of ENKD1 stabilizes microtubules and interferes with primary cilium formation

Given that ENKD1 localizes to microtubule-based structures in cells, we hypothesized that it might play important roles during centrosome and cilium biogenesis and function by regulating microtubules. To test this, we first examined the phenotypic consequences of its overexpression on microtubule polymerization, organization and stability in cells. In low-expressing cells, GFP-ENKD1 localized to the centrosome and the microtubule network was intact (Fig. 2A). However, its overexpression induced formation of microtubule bundles and disrupted the organization of microtubules. Likewise, overexpression of GFP-ENKD1(1-150) and GFP-ENKD1 (1-250) induced formation of microtubule bundles (Fig. 3A). Of note, the percentage of bundles in GFP-ENKD1 (1-150)-expressing cells were much lower than that of GFP-ENKD1 and GFP-ENKD1 (1-250)-expressing cells (Fig. S3B). Given their ability to bundle microtubules, we next tested whether GFP-ENKD1 and its N-terminal fragments bind to microtubules in cells. To this end, we performed *in vitro* microtubule sedimentation experiments with lysates prepared from cells transfected with GFP-ENKD1 (FL, 1-150 a.a., 1-250 a.a., 250-346 a.a.) or LAP and found that GFP-ENKD1 and its N-terminal fragments, but not LAP and GFP-ENKD1 (250-346 a.a.), co-pelleted with microtubules (Fig. 3B).

**Figure 3.**
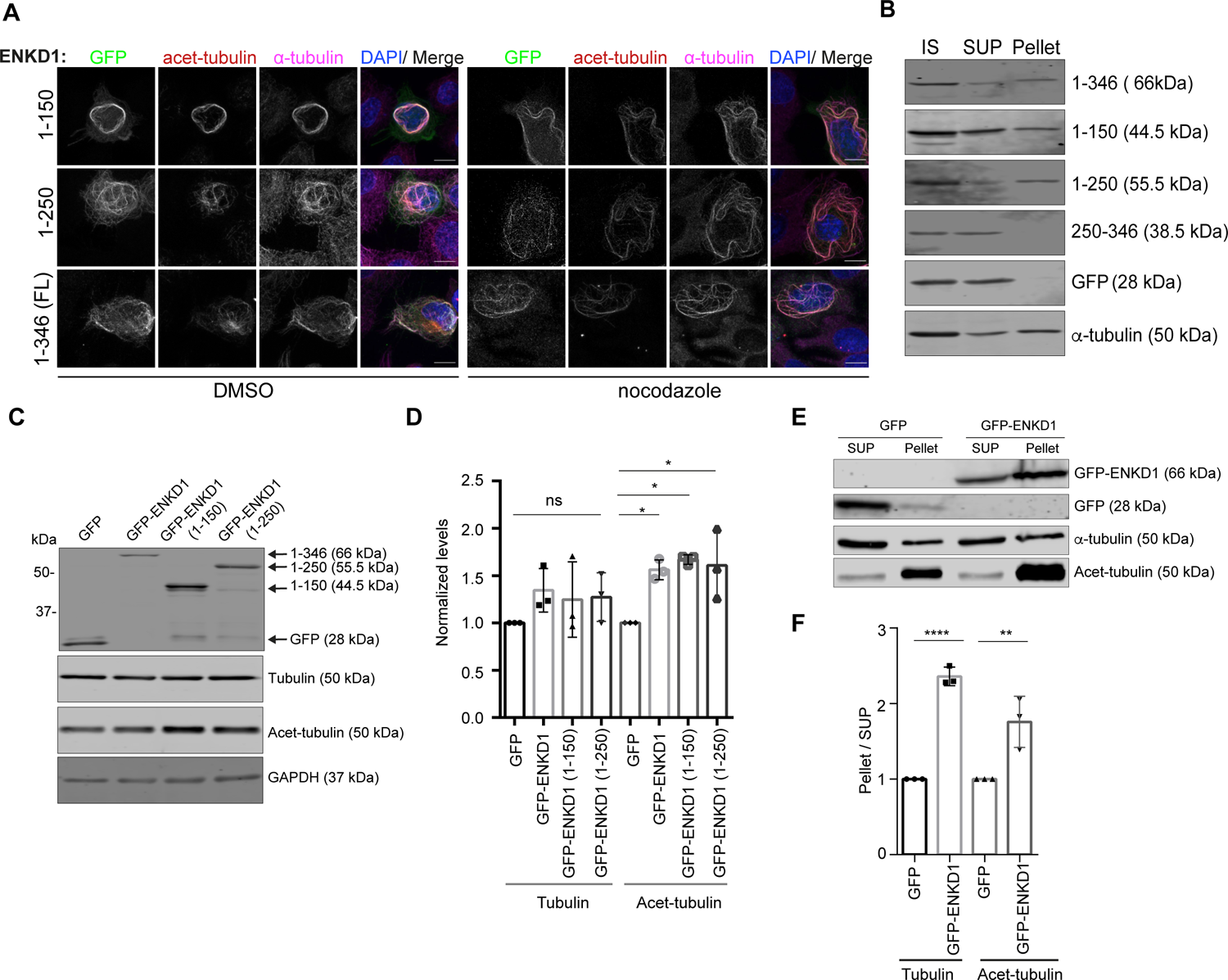
Overexpression of ENKD1 stabilizes microtubules and interferes with primary cilium assembly. (A) Localization of GFP-fusions of ENKD1 and its deletion mutants in transiently-transfected high-expressing RPE1 cells. RPE1cells were with the indicated constructs, treated with DMSO or nocodazole, fixed and stained for GFP, acetylated-tubulin and microtubules (α-tubulin). Scale bar: 10 `μ`m (B) Analysis of microtubule association of ENKD1 and its deletion mutants in cells. *In vitro* microtubule pelleting assays were performed using extracts from HEK293T cells transiently expressing GFP-tagged full-length ENKD1 and its deletion mutants. Initial sample (IS), supernatant/flowthrough (SUP), and pellet samples were spun down on glycerol cushion, separated by SDS-PAGE and immunoblotted with antibodies against GFP and tubulin. (C) Effect of overexpression of GFP-ENKD1 and its deletion constructs on total tubulin and acetylated tubulin levels. Lysates from HEK293T cells expressing the indicated proteins were separated by SDS-PAGE gel and immnoblotted using antibodies against GFP, α-tubulin, acetylated tubulin, and GAPDH. (D) Quantification of 3C. Band intensities were measured using ImageJ and normalized intensities were plotted on the graph. Data were derived from three experimental replicates. Mean tubulin intensity of GFP is 1.000; of GFP-ENKD1 is 1.344; of LAP ENKD1 (1-150) is 1.247; and of GFP-ENKD1 (1-250) is 1.272. Mean acetylated-tubulin intensity of GFP is 1.000; of GFP-ENKD1 is 1.562; of GFP-ENKD1 (1-150) is 1.668; and of GFP-ENKD1 (1-250) is 1.609. (E) Effect of GFP-ENKD1 expression on soluble and polymerized tubulin. Lysates from HEK293T cells expressing the indicated proteins were prepared in taxol-containing buffer. Supernatant (SUP) and pellet fractions were separated by centrifugation of samples at 12.000 x g for 10min at 4°C. Lysates were separated by SDS-PAGE gel and immunoblotted for GFP, α-tubulin, and acetylated tubulin. (F) Quantification of 3E. Band intensities were measured using ImageJ and polymerized to soluble pool ratios were calculated by dividing pellet band intensities to supernatant band intensities. Data were derived from three experimental replicates. Mean tubulin intensity of GFP is 1.000 and of GFP-ENKD1 is 2.359; mean acetylated-tubulin intensity of GFP is 1.000and of GFP-ENKD1 is 1.758.

Because these bundles co-localized with acetylated tubulin that marks stabilized microtubules, we next tested whether GFP-ENKD1 and its N-terminal fragments promoted microtubule stabilization. While microtubules depolymerized in control untransfected cells after nocodazole treatment, microtubule bundles persisted in cells expressing GFP-ENKD1 and its (1-150) and (1-250) a.a fragments (Fig. 3A). Consistent with their microtubule-stabilizing activity, immunoblotting of whole cell lysates from transfected cells showed that cells expressing GFP-ENKD1 and its N-terminal fragments resulted in an increase in acetylated tubulin levels as compared to control cells, while the total cellular abundance of alpha-tubulin remained unaltered (Fig. 3C, 3D).

Furthermore, we performed microtubule pelleting assay to compare the relative abundance of polymerized tubulin in control cells and cells expressing GFP-ENKD1. The fraction of alpha-tubulin and acetylated-tubulin found in the pellet that contains polymerized tubulin were higher as compared to the supernatant that contains soluble tubulin (Fig. 3E, 3F). These experiments together show that ENKD1 interacts with microtubules in cells and promotes microtubule polymerization and stabilization upon overexpression.

### ENKD1 binds to microtubules and promotes their polymerization *in vitro*

Having established the localization and interaction of ENKD1 with microtubules in cells, we next asked whether ENKD1 binds to microtubules directly. To this end, we first expressed recombinant maltose-binding protein (MBP)-tagged ENKD1 and MBP (negative control) in bacteria and purified them using amylose beads (Fig. S4A). We then performed *in vitro* microtubule sedimentation assays with bovine brain microtubules. MBP-ENKD1, but not MBP, co-pelleted with microtubules, which confirms a direct interaction between ENKD1 and microtubules (Fig. 4A, S4B). To calculate the binding dissociation constant (Kd) of this interaction, we performed sedimentation assays with a constant ENKD1 concentration (3 µM) and increasing microtubule concentrations (Fig. 4A: 0.01 to 1 µM; Fig. S5B: 0 to 4 µM). Additionally, we performed reciprocal experiments using a constant concentration of microtubules (1 µM) and increasing ENKD1 concentrations (1 to 4 µM) and verified the direct association of ENKD1 with microtubules (Fig. S4C). By plotting ENKD1 bound to microtubules versus microtubule concentration, we calculated the Kd of ENKD1 as 0.16 ± 0.08 µM (n=4).

**Figure 4:**
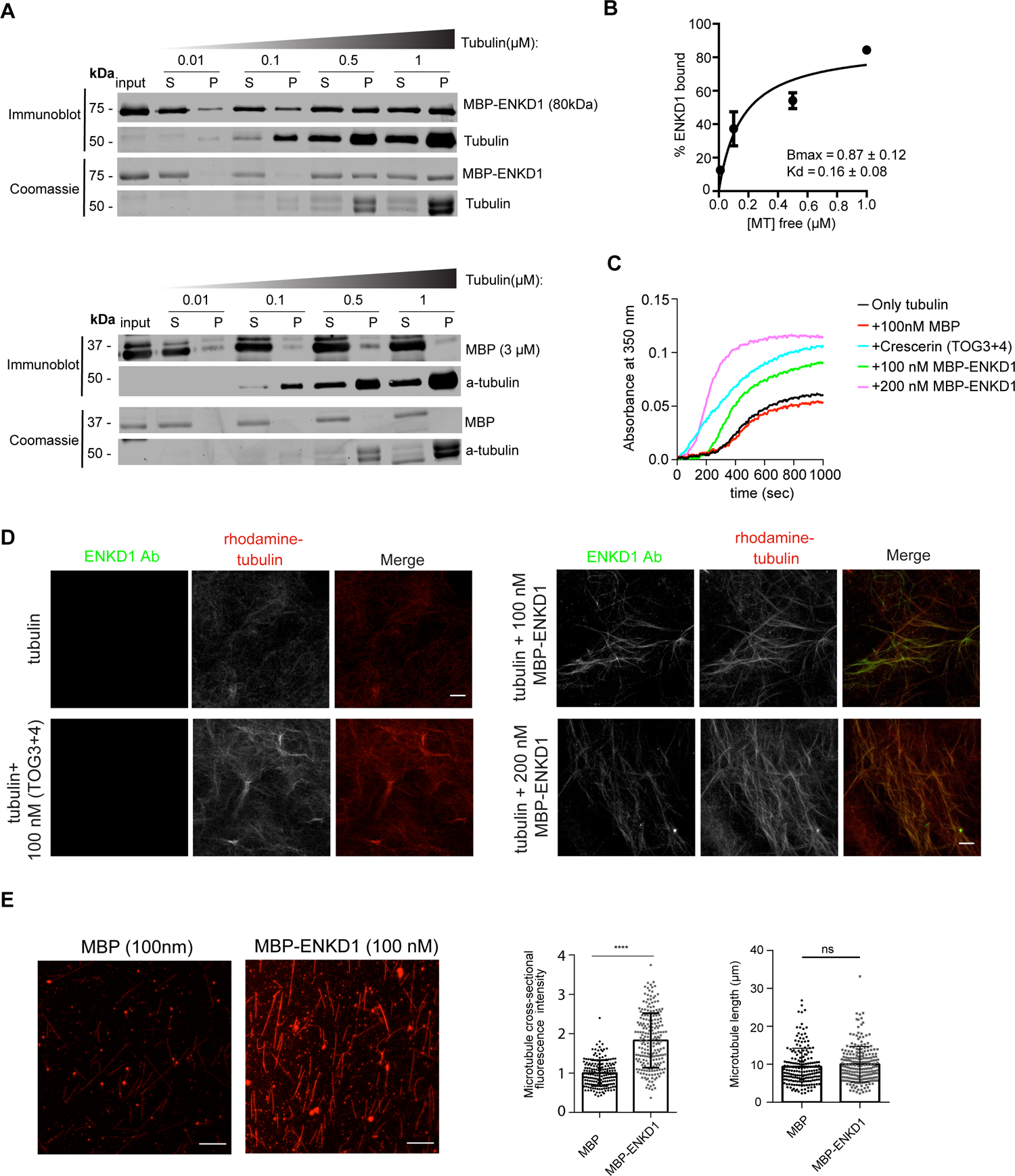
ENKD1 is a microtubule-associated protein that promotes microtubule polymerization. (A) *In vitro* microtubule cosedimentation assay using 3 μM of the indicated proteins and 0.01, 0.1, 0.5, and 1 μM Taxol-stabilized MTs. After incubation of proteins with MTs, reactions spun down (55,000 rpm for 10 min) on 40% glycerol cushion to pellet the polymerized MTs. Input, supernatant (S) and pellet (P) fractions were separated on SDS-PAGE gel and proteins were detected by Coomassie Blue staining and immunoblotting using antibodies against ENKD1, MBP and alpha-tubulin. (B) Quantification of ENKD1 microtubule binding affinity. Band intensities were quantified, background was subtracted and the dissociation constant (*K*d) was determined from the best-fit curve and calculated by using Prism 6 software (one site-specific binding). Data were derived from four experimental replicates. (C) *In vitro* MT polymerization assay was performed by measuring absorbance of the indicated reaactions at 350 nm. (D) Samples form 4C were fixed in glutaraldehyde and spun down on to coverslips for imaging. Samples were immunostained with an antibody against ENKD1. Microtubules were visualized by rhodamine tubulin. Scale bar:10μm (E) *In vitro* MT bundling assay was performed using 2 μM rhodamine-labeled taxol polymerized MTs. Cross-sectional fluorescence intensities were measured and average peak intensities were plotted for respective samples. Mean MT length were quantified by measuring individual lengths of microtubules in different samples. ns, not significant; ****P<0.0001.

Given that the Kd of established MAPs were reported as 1-2 µM, our results identify ENKD1 as a new MAP that binds to microtubules with high affinity (Cheeseman et al., 2006). Next, we examined whether and if so how ENKD1 regulates microtubule polymerization and organization *in vitro*. First, we used microscopy- and turbidity-based assays to examine whether ENKD1 binds to microtubules and promotes polymerization *in vitro*. For the turbidity assays, we mixed 100 nM and 200 nM MBP-ENKD1 with tubulin and quantified tubulin polymerization at 37°C by monitoring light scattering at 350 nm.

As controls, MBP (negative control) did not promote microtubule polymerization and TOG3-4 domains of Crescerin/TOGARAM1 (positive control) increased the rate of microtubule polymerization *in vitro* (Das et al., 2015). Increasing concentrations of ENKD1 reduced the lag time of polymerization and increased the microtubule mass at steady state (Fig. 4C). To visualize the microtubules network induced by MBP-ENKD1 and to test whether ENKD1 binds to this network, we spun down the polymerization reactions containing rhodamine-tubulin on coverslips and stained them with ENKD1 antibody. In agreement with the turbidity assays, microscopic analysis of the reactions showed that addition of MBP-ENKD1 to tubulin increased the polymer mass relative to tubulin alone. Importantly, ENKD1 co-localized with the microtubules, further validating its direct microtubule association (Fig. 4D).

Finally, we examined whether ENKD1 bundles taxol-stabilized, rhodamine-labeled microtubules by microscopic analysis (Zhu et al., 2018). Microtubules remained short and thin in samples containing MBP, which was used as a negative control.

Addition of 100 nM MBP-ENKD1 to the reaction resulted in an increased microtubule mass as compared to the MBP control. We quantified microtubule-crosslinking activity and microtubule length as previously described (Zhu et al., 2018). While microtubule length did not change between MBP and MBP-ENKD1-containing samples, the microtubule cross-sectional intensity was higher in the presence of MBP-ENKD1 (Fig. 4E). Collectively, these results indicate that ENKD1 regulates microtubule dynamics by promoting microtubule polymerization.

### ENKD1 has extensive proximity interactions with centrosome and microtubule-associated proteins

To gain insight into cellular functions of ENKD1, we identified its proximity interaction map using the BioID approach. Using lentiviral transduction, we generated HEK293T cell line that stably express N-terminal V5-BirA(R118G) (hereafter BirA*) fusion of ENKD1. Incubation of the stable line biotin followed by staining for the biotinylated proteins and centrosome marker demonstrated that BirA*-ENKD1 localized to and induced biotinylation at the centrosome, as assessed by staining for streptavidin and gamma-tubulin (Fig. 5A). Immunoblotting of lysates from the stable line and the eluate after the streptavidin pulldown validated the expression of BirA*-ENKD1 as well as its pulldown along with the biotinylated proteins in its proximity (Fig. 5B).

**Figure 5.**
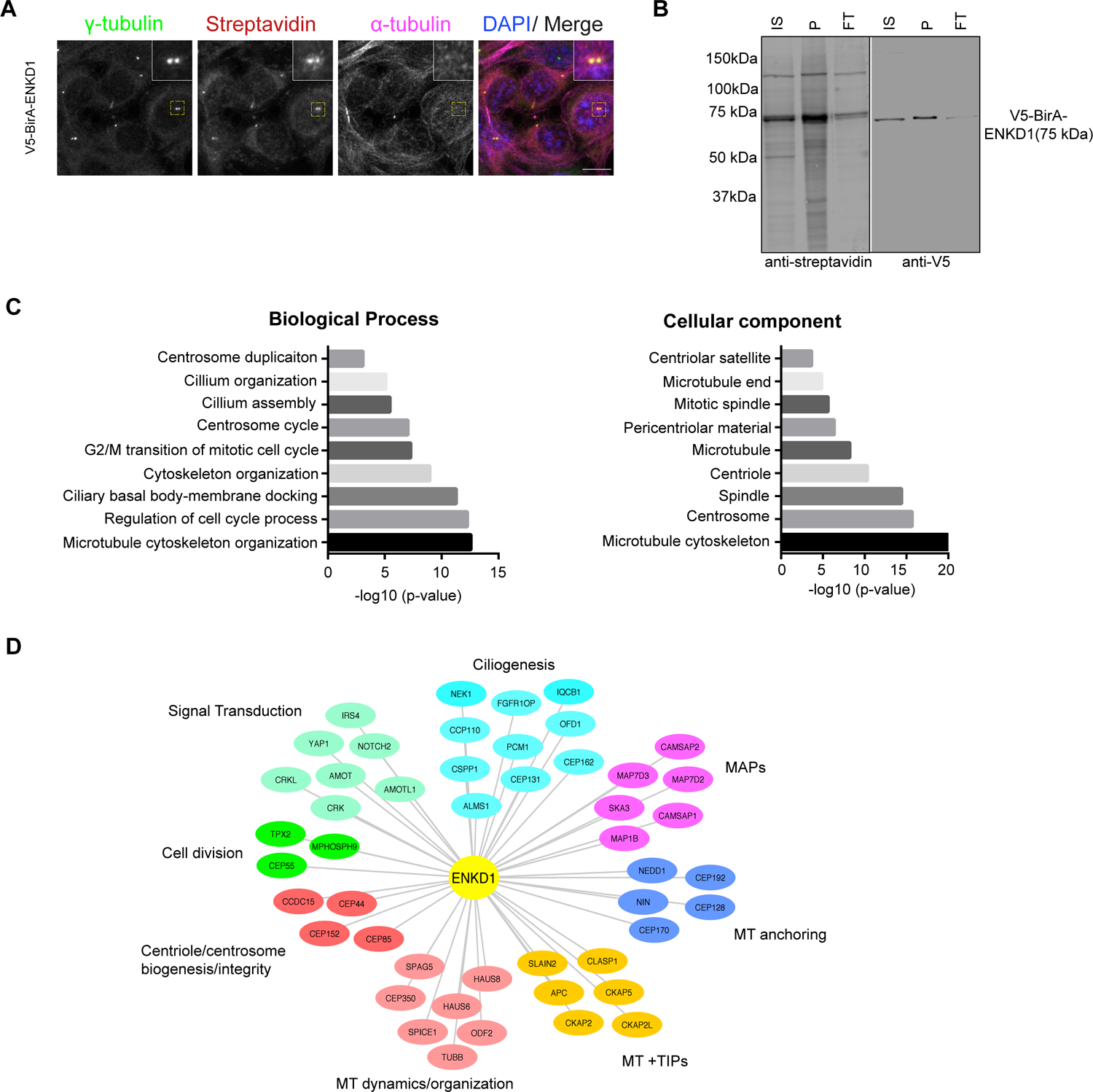
ENKD1 has extensive proximity interactions with centrosome/cilium proteins and MAPs. (A) Validation of V5-BirA*-ENKD1-induced biotinylation by immunofluorescence Localization of V5-BirA*-ENKD1 relative to markers of the centrosome and microtubules. HEK293T::V5-BirA*-ENKD1 cells were incubated with 50 `μ`M biotin for 18 h and immunostained with fluorescent streptavidin and antibodies against `γ`-tubulin and α-tubulin. Scale bar, 10 `μ`m. (B) Validation of V5-BirA*-ENKD1-induced biotinylation and pulldown of biotinylated proteins by immunoblotting. Cells were lysed, and biotinylated proteins were precipitated by streptavidin pull down. The initial sample (IS), immunoprecipitated biotinylated proteins (P), and flow-through (FT) were run on a gel and immunoblotted with antibodies against streptavidin and V5. (C) GO-enrichment analysis of the ENKD1 proximity interactors based on their biological process and cellular compartment. The x-axis represents the log-transformed p-value (Fisher’s exact test) of GO terms. (D) Interaction networks of ENKD1 based on biological process and cellular compartment. The ENKD1 proximity interactors were grouped using g-Profiler functional annotation tool. The interaction networks visualized for the biological processes include microtubule organization/dynamics, microtubule anchoring, MT+TIPs, MAPs, primary cilium biogenesis, cell division, signal transduction, and centriole biogenesis/integrity.

For mass spectrometry experiments, we performed large scale streptavidin pulldowns from BirA*-ENKD1-expressing cells incubated with 50 µM biotin for 18 h. As a control, we used the mass spectrometry data derived from streptavidin pulldown from bitoin-induced HEK293T::V5-BirA* cells (Arslanhan et al., 2021). We defined the high confidence ENKD1 interactome by applying two different filters to the mass spectrometry data derived from two experimental replicates. First, we analyzed the data by significance analysis of interactome (SAINT), which assigns confidence scores to each bait-prey interaction (Teo et al., 2014). To filter out low confidence and non-specific interactions, we used Bayesian false discovery rare (BFDR) cutoff of <0.03. Second, we only accounted for proteins identified with spectral counts greater than 2 in each experimental replicate. These analyses together yielded 204 proteins as high-confidence ENKD1 proximity interactors.

To generate hypothesis about ENKD1 cellular functions, we performed Gene Ontology (GO) analysis to determine which biological processes and cellular compartments are enriched among 204 high confidence ENK1 proximity interactors. Consistent with its cellular localization, ENKD1 had abundant proximity interactions with centrosome proteins, centriolar satellites and MAPs (Fig. 5C). GO analysis for biological processes revealed functions associated with these compartments including the highly enriched “microtubule organization”, “cell cycle”, “cilium assembly” and “centrosome duplication” categories (Fig. 5C). In agreement, ENKD1 was recently characterized for its functions during proliferation, migration and invasion of non-small cell lung cancer cells (Song et al., 2021). Together with cellular localization and overexpression phenotypes of ENKD1, we aimed to use its proximity interactome to generate hypothesis on the specific centrosome-, cilium- and microtubule-associated processes regulated by ENKD1. To this end, we classified the related proteins to specific functions and compartments and generated a sub-interaction network for ENKD1 (Fig. 5D). In agreement with its cellular localization, MAPs and centrosome/satellite proteins involved in cell cycle progression, centrosome duplication, centriole integrity and cilium biogenesis were enriched in the ENKD1 proximity interactome. Overall, the proximity map of ENKD1 provides further support for its functions at the centrosome, the cilia and/or microtubule-associated processes such as cell division.

### ENKD1 is required for cilium length and ciliary content regulation

Cellular localization, interactions, and evolutionary conservation of ENKD1 suggests potential functions during centrosome, cilium and/or microtubule mediated processes. In order to elucidate the functions and mechanisms of MAPs at the primary cilium, we specifically examined whether and if so how ENKD1 regulates primary cilium assembly and function. To this end, we assayed ENKD1-depleted cells using a set of different functional assays. For ENKD1 depletion, we infected IMCD3 cells with lentivirus expressing a short hairpin RNA (shRNA) targeting mouse ENKD1 or control shRNA, selected cells with puromycin and generated cell lines stably expressing control and ENKD1 shRNAs. Immunoblotting and qPCR analysis confirmed that expression of ENKD1 shRNA results in its efficient depletion (Fig. S5A).

ENKD1-depleted cells ciliated at similar percentages like control cells (Fig. 6A, 6B). Notably, the cilia that formed in ENKD1-depleted cells were shorter as compared to control cells (control: 3.1 ±0.03 µm, shENKD1: 2.8±0.03 µm), which identifies ENKD1 as a regulator of cilium length (Fig. 6A, 6C). The bidirectional transport of cargoes (i.e. tubulin) by the intraflagellar transport (IFT)-A and IFT-B machineries is one of the best characterized mechanisms described for cilium length regulation (Broekhuis et al., 2013; Wang et al., 2021). To test whether defective ciliary recruitment of IFT to the cilium underlies the cilium length regulation by ENKD1, we quantified the ciliary concentration of anterograde IFT-B component IFT88. Ciliary IFT88 concentration was reduced in ENKD1-depleted cells, suggesting that reduced intraflagellar transport might underlie their cilium shortening phenotype (Fig. 6D).

**Figure 6.**
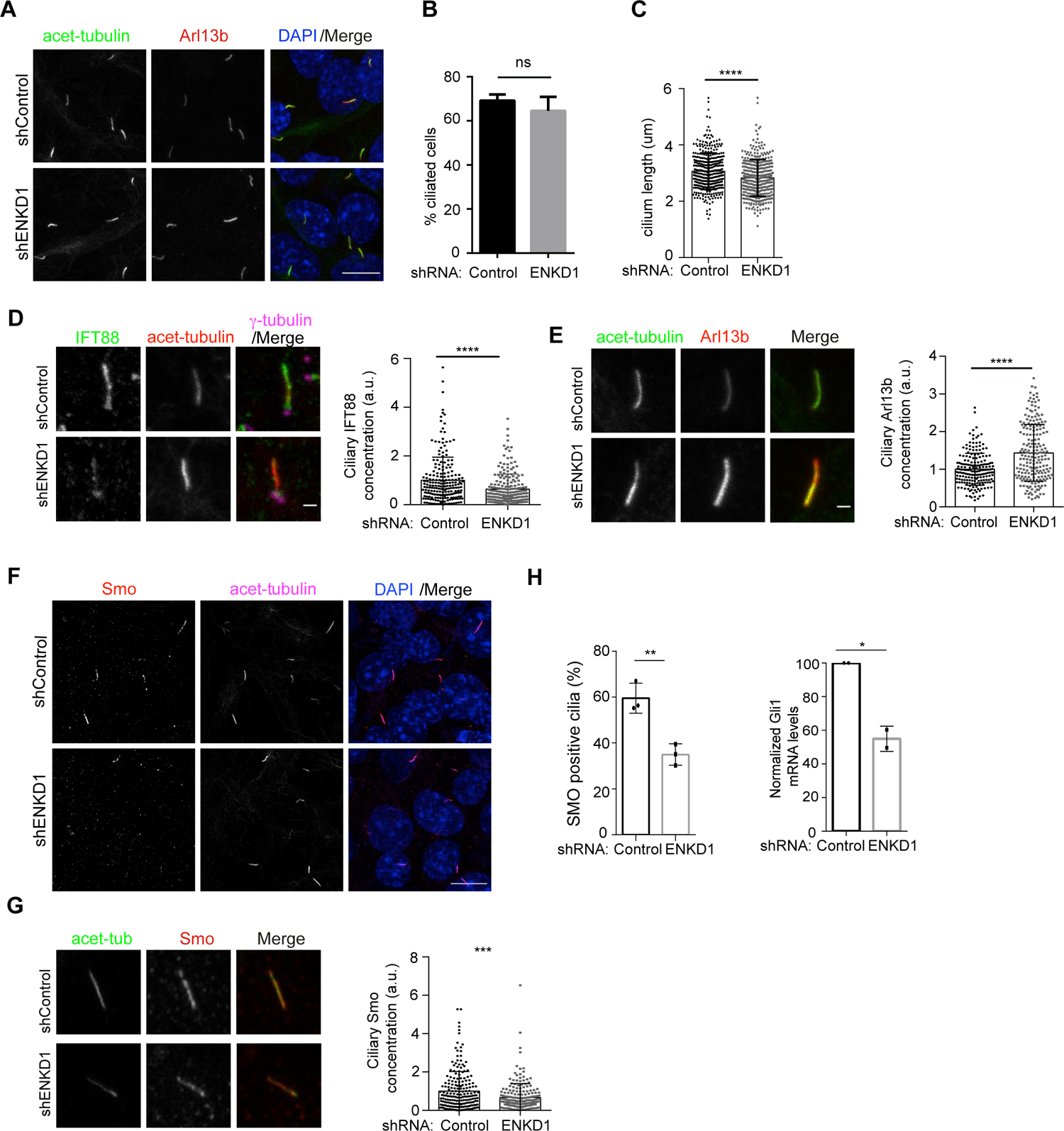
ENKD1 is required for regulation of cilium length, ciliary content and Hedgehog signaling. (A) Effect of ENKD1 depletion on cilium formation. Control shRNA and IMCD3 shENKD1-expressing stable cells were serum-starved for the 24 h, and percentage of ciliated cells was determined by staining cells for acetylated tubulin, Arl13b, and DAPI. Scale bar, 10 `μ`m (B) Quantification of ciliation percentage as represented in 6A. Results shown are the mean of three independent experiments. % ciliated cells in shControl is 69.14 (±1.646); % ciliated cells in shControl is 64.57 (±3.643). There is no significant difference between control and IMCD3 shENKD1 cells (t-test). (C) Quantification of cilium length as represented in 6A. Results shown are the mean of three independent experiments. Mean cilium length in shControl is 3.067 `μ`m (±0.03303, N=380); mean cilium length in shENKD1 is 2.831 `μ`m (±0.03426, N=367) from three independent experiments. (D) Effects of ENKD1 depletion on the ciliary IFT88 concentration. Control shRNA and shENKD1-expressing IMCD3 cells were serum-starved for 24 h, fixed and stained for IFT88, acetylated tubulin and `γ`-tubulin. Mean IFT88 concentration in shControl cells is 1.000 (±0.06531, N=213); in shENKD1 cells is 0.6390 (±0.04145, N=217). (E) Effects of ENKD1 depletion on the ciliary Arl13b concentration. shControl and shENKD1 IMCD3 cells were serum-starved for 24 h, fixed and stained for acetylated tubulin and Arl13b. Mean Arl13b concentration in shControl cells is 1.000 (±0.03051, N=195); in shENKD1 cells is 1.435 (±0.05299, N=203). (F) Effect of ENKD1 depletion on ciliary recruitment of Smoothened (SMO). shControl and shENKD1 expressing IMCD3 cells were serum-starved for 24 h, treated with 200 nM SAG for 4h, fixed and stained for SMO, acetylated tubulin (Ac. tub), and DAPI. Percentage of Smo-positive cilia was quantified. % SMO localized to cilia in shControl cells is 59.52; in shENKD1 cells 34.94.(n=3). Effect of ENKD1 depletion on Gli1 mRNA levels were quantified by qRT-PCR. Normalized mean Gli1 mRNA levels in shControl cells is 100; in shENKD1 cells is 54.9725. p= 0.013. (G) Quantification of the ciliary concentration of SMO from two independent experiments. shControl and shENKD1 expressing IMCD3 cells were serum-starved for 24 h, treated with 200 nM SAG for 4h, fixed and stained for SMO, acetylated tubulin (Ac. tub), and DAPI. Mean SMO concentration in shControl cells is 1.000 (± 0.7091, N=207); in shENKD1 cells is 0.6731 (± 0.05021, N=207) Scale bar, 10 `μ`m

Proper structure and length of the primary cilium is required for regulation of the cilium composition (Broekhuis et al., 2013; Keeling et al., 2016). To determine whether ENKD1 regulates ciliary levels of signaling proteins, we quantified the ciliary concentration of the small GTPase Arl13b, which binds to the ciliary membrane via palmitoylation and is required for ciliary trafficking of Hedgehog proteins (Caspary et al., 2007; Gigante et al., 2020). ENKD1-depleted cells had higher levels of ciliary Arl13b compared to control cells (Fig. 6E), which indicates deregulation of ciliary content upon ENKD1 loss. In addition to the ciliary proteins that constitutively localize to the primary cilium, we assayed ENKD1-depleted cells for their ability to respond to Hedgehog ligands. During pathway activation, Smo accumulates in the primary cilium and initiates a cascade that results in transcriptional activation of Hedgehog target genes (Wheway et al., 2018). Upon stimulation of IMCD3 cells with Smoothened agonist (SAG) for 4 h, there was a 0.58-fold reduction in the percentage of Smo-positive cilia in ENKD1-depleted cells relative to control cells (Fig. 6H). In agreement, ciliary Smo concentration was also reduced upon ENKD1 depletion. Ciliary Smo recruitment defect was rescued by stable expression of mNG-fusions of human full-length ENKD1 and its N-terminal (1-250 a.a.) and C-terminal (250-346) domains, confirming its specificity to ENKD1 depletion and highlighting the cooperative functions of N- and C-terminal domains during Hedgehog response (Fig. S6B). Finally, we examined the transcriptional response of ENKD1-depleted cells to Hedgehog pathway activation. As compared to control cells that had robust activation of Gli1 expression after 24 h SAG treatment, ENKD1-depleted cells failed to upregulate their expression (Fig. 6H). Collectively, these results identify ENKD1 as a regulator of ciliary length, trafficking and signaling.

## Discussion

In this study, we identified ENKD1 as a new MAP that regulates microtubule polymerization and organization in vitro and in cells. We showed that ENKD1 localizes to the centriole wall and the ciliary axoneme, and functions during cilium assembly, Hedgehog signaling and ciliary content regulation. These results advance our understanding of how centriolar and ciliary MAPs contribute to building and maintaining functional primary cilia competent for Hedgehog signaling.

Although several ciliary and centriolar MAPs were identified and characterized, the parts list for the centriolar and ciliary MAPs is incomplete (Conkar and Firat-Karalar, 2020; Mirvis et al., 2018). This hampers progress in uncovering the molecular mechanisms by which the length, stability and architecture of centrioles and axonemes are built from microtubules and associated proteins. Our results contribute to addressing these questions in two ways. First, we defined ENKD1 as a centriolar and ciliary MAP. Second, the proximity map of ENKD1 identified its proximity relationships with other MAPs and thus, it provides a resource for future studies aimed as identifying centriolar and ciliary MAPs. To predict the ciliary interactors of ENKD1 in its proximity map, we compared it to the recently published primary cilium proximity proteome and found several MAPs as overlapping proteins (CKAP2L, SKA3, CKAP2, CSPP1, MAP4, NIN) (May et al., 2021). Analogous to the cytoplasmic MAPs that function as part of large protein complexes, ENKD1 might also form complexes with other ciliary proteins to regulate microtubule dynamics during cilium assembly and maintenance. A number of ENKD1 proximity interactors stand out by their previously described ciliary functions.

CEP162 was reported as an axoneme-recognition protein that mediates the interaction between transition zone components and the axoneme (Wang et al., 2013). The Joubert-associated CSPP1 was shown to localize to the ciliary axoneme/tip and regulate cilium length and Hedgehog signaling (Frikstad et al., 2019). Future studies are required to define the physical interactors of ENKD1 at the centrioles and cilia, which will provide insight into how ENKD1 regulates axonemal microtubule dynamics and stability and the mechanism by which it mediates its ciliary functions.

We investigated the roles of the different ENKD1 fragments in its localization and function and showed that ENKD1 localizes to the centrosome and primary cilium through distinct regions, which enabled us to study the relationship between its localization and function. The axonemal localization of ENKD1 depends on its ability to interact with microtubules, further supporting its functions as a ciliary MAP. Notably, the length of the ENKD1-positive axoneme relative to the cilium length substantially varied from cilium to cilium, which was previously reported for components of the Inversin compartment of primary cilium (Bennett et al., 2020). Because ENKD1 did not localize with inversin, this result suggests that ENKD1 association with the ciliary axoneme might require specific tubulin modifications or ciliary binding partners. Addressing these possibilities in future studies might potentially identify a new sub-compartment within the primary cilium.

As for the centrosomal localization of ENKD1, it is mediated by its N-terminal (1-250 a.a.) microtubule-binding fragment and the C-terminal enkurin domain (250-346 a.a.). Intriguingly, expansion microscopy revealed distinct pools of ENKD1 at the centriole wall and PCM. Given that only N-terminal domain of ENKD1 binds to microtubules, it is possible that ENKD1 is targeted to the centriole wall by this region and to the PCM by its enkurin domain. Identification of the high-resolution localization and interactors of ENKD1 (1-250) and ENKD1 (250-346) in future studies will address these possibilities. We also highlight that the C-terminal enkurin domain of ENKD1 is also present in another ciliary protein named ENKUR, which was reported as a regulator of motile cilia assembly in zebrafish and tracheal cultures (Sigg et al., 2017). Moreover, enkurin domain of ENKUR was shown to mediate its interaction with transient receptor potential (TRP) channels and calmodulin in sperm (Sutton et al., 2004). While future studies are required to investigate whether ENKD1 also functions during motile cilia assembly, the specific activities and interactions mediated by the enkurin domain of ENKD1 in mammalian cells is not known and whether it also function during motile cilia assembly needs to be investigated.

A key finding of our study is the identification of ENKD1 as a new regulator of microtubule polymerization, organization and bundling. *In vitro* microtubule co-sedimentation experiments showed that ENKD1 binds to microtubules with high affinity. Given that microtubules are negatively charged on their surface and a subset of microtubule-binding regions were characterized as positively-charged (i.e. CSAP) (Bompard et al., 2018). Although this high affinity can be explained by the enrichment of ENKD1 in basic amino acid residues throughout the protein (pI =10.46), it does not explain why N-terminus targets ENKD1 to the microtubules. In contrast to its microtubule affinity, the mechanism by which ENKD1 promotes microtubule polymerization cannot be predicted from domain composition and organization. ENKD1 does not have conserved domains (i.e. TOG) involved in enabling proteins to function as polymerases (Farmer and Zanic, 2021). Structural characterization of the interaction between ENKD1 and microtubules will precisely map the microtubule-interaction surface of ENKD1 and possibly explain how ENKD1 binds to and regulates microtubules.

Functional assays revealed that microtubule-associated activities of ENKD1 are required for primary cilium assembly and ciliary content, particularly in response to Hedgehog ligands. We envision two possible mechanisms by which ENKD1 is involved in Hedgehog response. Given its microtubule-associated functions, impaired Hedgehog signaling in ENKD1-depleted cells might be an indirect consequence of architectural defects of the axoneme and the resulting ciliary trafficking defects. In agreement, ciliary recruitment of the IFT-B complex was defective upon ENKD1 loss. Additionally, ENKD1 might regulate Hedgehog signaling through its enkurin domain, which mediates interaction of ENKUR with TRP channels (Sutton et al., 2004). Using evolutionary rpoteomics, Sigg et al. showed that ENKD1 was clustered with TRP-channel related ciliary proteins such as ENKUR, PKD1 and PKD2 (Sigg et al., 2017). They also discovered that ciliary regulators of Hedgehog receptor pathways were linked to cilia before the origin of bilateria and TRP channels before the origin of animals, supporting evolutionarily conserved functions for ENKD1 ciliary signaling. Future investigation of ENKD1 functions in different mammalian cell types and ciliated organisms will unravel the full extent for ENKD1 functions at the primary cilium and motile cilia.

Given that ENKD1 regulates microtubule polymerization and organization *in vitro*, the lack of ciliogenesis defect in ENKD1-depleted cells were surprising. Unlike ENKD1, previously described centriolar and ciliary MAPs involved in microtubule polymerization and/or stability (i.e. CSPP1, Crescerin/TOGARAM1) were identified as critical regulators of axoneme length (Dacheux et al., 2015; Das et al., 2015; Frikstad et al., 2019; Latour et al., 2020). We envision two possibilities for why ENKD1 depletion did not interfere with ciliogenesis. First, there might be functional redundancy between ENKD1 and other ciliary MAPs. Second, ENKD1 might be required for the the architectural integrity of centrioles and the axoneme, which could have only been identified by structural studies (i.e. cryo-EM). The second model is exemplified for CEP104 and Crescerin, both of which are ciliary MAPs reported to determine the geometry of the distal axoneme.

Structural studies in *Tetrahymena* mutants identified Crescerin as a regulator of B-tubule length and CEP104 as a regulator of A-tubule elongation (Louka et al., 2018). In future, these models should be tested by identification of ENKD1 interactors and high-resolution imaging of centrioles and cilia in ENKD1-depleted cells using cryo-EM.

Based on our data, we propose several mechanisms by which ENKD1 regulates cilium biogenesis as a MAP. First, ENKD1 might contribute to the stabilization of the centriolar and axonemal microtubules, which are highly modified and stable. Given that ENKD1 overexpression increases microtubule acetylation, ENKD1 might regulate the ciliary axoneme by regulating the stability of axonemal microtubules, either directly or indirectly through cytoplasmic microtubules. Second, ENKD1 might be a component of the machinery that controls cilium length. At the cilia, it might cooperate or compete with other ciliary MAPs involved in axonemal microtubule polymerization and depolymerization. For example, Crescerin and CEP104 promote microtubule polymerization *in vitro* via their conserved tubulin-binding TOG domain, which were shown to be required for axoneme elongation in mammalian cells and *C. elegans*, respectively (Das et al., 2015; Frikstad et al., 2019; Latour et al., 2020; Satish Tammana et al., 2013; Yamazoe et al., 2020). Kif7 negatively regulates axoneme length by reducing the rate of microtubule growth and increasing the frequency of microtubule catastrophe (He et al., 2014; Jiang et al., 2019). ENKD1 might directly or indirectly regulate rescue and catastrophe events at the plus tips of the ciliary axoneme. TIRF-based imaging of the *in vitro* reconstitution experiments between ENKD1 and dynamic microtubules might further contribute to our understanding of the mechanisms by which ENKD1 regulates axonemal microtubules.

## Materials and Methods

### Plasmids

Full-length cDNAs for human ENKD1 (GenBank accession no. NM_ 032140.3) and its deletion mutants were amplified by PCR and cloned into pDONR221 plasmid using Gateway technology (Life Sciences). Their N-terminal LAP (GFP-Tev-S-tag)-fusions or His-MBP (Maltose-binding protein)-fusions were generated by Gateway cloning into pEF-FRT-LapC DEST (provided by Max Nachury, UCSF) or pDEST-HisMBP, respectively. pCDH-EF1-mNeonGreen-T2A-Puro plasmid was used to generate mNG-ENKD1, mNG-ENKD1 (1-250) and mNG-ENKD1 (250-246) expression vectors. *Homo sapiens* Inversin and IFT88 were cloned into pCDH-EF1-mScarlet-T2A-Puro plasmid generate their mScarlet-fusions. pDONR221 plasmid containing cDNA of human TOGARAM1 (GenBank accession no. NM_001308120.2) was a kind gift from R. Roepman. DNA fragment spanning the TOG3 and TOG4 domains of human TOGARAM1 (amino acids 1240-1773) was PCR amplified and subsequently cloned into pDEST-HisMBP vector for bacterial expression. Mouse ENKD1 shRNA (nucleotides targeting 403-423 bp, 5′-CGCTCACCCAAGTATGACAAT-3′) were cloned into pLKO.1 (Invitrogen).

### Cell culture, transfection and lentivirus

Human telomerase immortalized retinal pigment epithelium cells (hTERT-RPE, ATCC, CRL-4000) and IMCD3 cells were cultured with Dulbecco’s modified Eagle’s Medium DMEM/F12 50/50 medium (Pan Biotech) supplemented with 10% Fetal Bovine Serum (FBS, Life Technologies) and 1% penicillin-streptomycin (Gibco). Human embryonic kidney (HEK293T, ATCC, CRL-3216) cells were cultured with DMEM medium (Pan Biotech) supplemented with 10% FBS and 1% penicillin-streptomycin. All cell lines were tested for mycoplasma by MycoAlert Mycoplasma Detection Kit (Lonza). RPE1 cells were transfected with the plasmids using Lipofectamine LTX according to the manufacturer’s instructions (Life Technologies). HEK293T cells were transfected with the plasmids using 1 μg/μl polyethylenimine, MW 25 kDa (PEI, Sigma-Aldrich, St. Louis, MO). For serum starvation experiments, cells were washed twice with PBS and incubated with DMEM/F12 50/50 supplemented with 0.5% FBS and 1% penicillin-streptomycin for the indicated times. For microtubule depolymerization experiments, cells were treated with 5 μg/ml nocodazole (Sigma-Aldrich) for 1 hour at 37°C.

Recombinant lentivirus expressing mNG-ENKD1, mNG-ENKD1 (1-250), mNG-ENKD1 (250-346), control and ENKD1 shRNAs were made by cotransfection of HEK293T cells with the respective transfer vectors, and packaging and envelope vectors. 48 h after transfection, supernatant was harvested, concentrated using Lenti-X Concentrator (Takara Bio Europe) and titered on IMCD3 cells using GFP-expressing lentivirus. For transduction, IMCD3 cells were plated in 24-well tissue culture plates, sequentially infected with an approximate MOI of 1 over two days. For short-term depletion, cells were assayed 6-10 days after initial infection. For long-term depletion, infected cells were selected with medium containing 3 μg/ml puromycin medium (Invivogen, CA) for 4 days and resulting heterogenous pools were used for functional assays. Depletion of ENKD1 was confirmed by immunoblotting and qPCR. To generate IMCD3 and RPE1 cell lines that stably express mNG-ENKD1, mScarlet-Inversin and/or mScarlet-IFT88, cells were infected with lentivirus expressing the fusion proteins, selected with medium containing 3 μg/ml (for IMCD3s), 10 μg/ml (for RPE1s) puromycin. Clonal cell lines were generated by serial dilutions of the heterogenous pool and screened for expression levels using immunoblotting and immunofluorescence.

### Antibodies

Primary antibodies used for immunofluorescence were rabbit anti ENKD1 (HPA041163, Sigma) at 1:200, mouse anti Arl13b (75-287, Neuromab) at 1:500, mouse anti acetylated tubulin (clone 6-11B, 32270, Thermo Fischer) at 1:10000, mouse anti gamma tubulin (Sigma, clone GTU-88, T5326) at 1:2000, mouse anti GFP (3E6) 1:750, mouse anti alpha tubulin (Sigma, DM1A) at 1:10000, mouse anti polyglutamylated-tubulin (AG-20B-0020, Adipogen) at 1:500), mouse anti C-NAP1 (Santa Cruz Biotechnology), mouse anti Centrin 3 (H0007070-MO1, Abnova), and rabbit anti IFT88 (139671-AP, ProteinTech). Secondary antibodies used for immunofluorescence experiments were AlexaFluor 488-, 568- or 633-coupled (Life Technologies) and Streptavidin 568. They were used at 1:2000. Anti-ENKD1 antibody was produced by immunizing rats with purified, bacterially expressed MBP fusion of full-length human ENKD1 (Koc University Animal Facility). ENKD1 antibody was affinity purified from rat anti-sera on nitrocellulose blots with purified, GST fusion of full length ENKD1.

Primary antibodies used for immunoblotting were rabbit anti ENKD1 (HPA041163, Sigma) at 1:1000, mouse anti alpha tubulin (Sigma, DM1A) at 1:10000, mouse anti acetylated tubulin (clone 6-11B, 32270, Thermo Fischer) at 1:10000, mouse anti beta-actin (Proteintech, 60008-1-Ig) at 1:10000, mouse anti alpha-tubulin (Sigma, DM1A) at 1:5000, mouse anti vinculin (sc-55465, Santa Cruz Biotechnologies) at 1:1000.

Secondary antibodies used for western blotting experiments were IRDye680- and IRDye 800-coupled and were used at 1:15000 (LI-COR Biosciences).

### Immunofluorescence, antibodies and microscopy

Cells were grown on coverslips, washed twice with PBS and fixed in either ice cold methanol at −20°C for 10 minutes or 4% PFA in cytoskeletal buffer (10 mM PIPES, 3 mM MgCl_2_, 100 mM NaCl, 300 mM sucrose, pH 6.9) supplemented with 5 mM EGTA and 0.1% Triton X for 15 min at 37°C (Hua & Ferland, 2017). After rehydration in PBS, cells were blocked with 3% BSA (Capricorn Scientific, Cat. # BSA-1T) in PBS + 0.1% Triton X-100 followed by incubation with primary antibodies in blocking solution for 1 hour at room temperature. Cells were washed three times with PBS and incubated with secondary antibodies and DAPI (Thermo Scientific, cat#D1306) at 1:5000 for 45 minutes at room temperature. Following three washes with PBS, cells were mounted using Mowiol mounting medium containing N-propyl gallate (Sigma-Aldrich).

Images were acquired with Leica DMi8 inverted fluorescent microscope with a stack size of 8 μm and step size of 0.5 μm in 1024×1024 format using HC PL APO CS2 63x 1.4 NA oil objective. Higher resolution images were taken by using HC PL APO CS2 63x 1.4 NA oil objective with Leica SP8 confocal microscope. Time lapse live imaging was performed with Leica SP8 confocal microscope equipped with an incubation chamber. For cell cycle and ciliation experiments, cells were imaged at 37°C with 5% CO_2_ imaged overnight with a frequency of 5-10 minutes per frame with 0.7 µm step size and 3.5 µm stack size in 512×512 pixel format.

All centrosomal and ciliary level quantifications were done in Image J (Schneider et al., 2012). Region of interest (ROI) around the centrosome or the primary cilium was determined using markers of the centrosome (gamma-tubulin) or the primary cilium (Arl13b, Acetylated tubulin) and the corresponding signals were quantified.

Cytoplasmic signal was subtracted from this value for every single cell. Statistical analysis was done by normalizing these values to their mean. Ciliary length was measured using acetylated tubulin or Arl13b as the ciliary length markers. Ciliary protein concentration was determined by dividing ciliary protein signal by ciliary length. All values were normalized relative to the mean of the overall quantification (=1).

### Fluorescence Recovery After Photobleaching

FRAP experiments were performed with Leica SP8 confocal microscope using FRAP module. Cells were incubated with 10% FBS in DMEM-12 and kept at 37°C with 5% CO_2_. ROI was set to 2.5 µm^2^ for centrosomal FRAP experiments. Since primary cilium length varied from cell to cell, a specific ROI was defined for each primary cilium. A z-stack of 4 µm with 0.5 µm step size was taken during pre and post bleaching for both centrosome and cilium FRAP experiments. Bleaching was done 2 iterations with 488 Argon laser with 100% power. Maximal projection of the files was performed in Leica LAS X software and analysis was done in ImageJ. Recovery graph quantifications, t-half and mobile pool quantifications were done using the equations as described (Sprague et al., 2004).

### Cell lysis and immunoblotting

Cells were lysed in 50 mM Tris (pH 7.6), 150 mM NaCI, 1% Triton X-100 and protease inhibitors for 30 min at 4°C followed by centrifugation at 15.000 g for 15 min. The protein concentration of the resulting supernatants was determined with the Bradford solution (Bio-Rad Laboratories, CA, USA). For immunoblotting, equal quantities of cell extracts were resolved on SDS-PAGE gels, transferred onto nitrocellulose membranes, blocked with TBST in 5% milk for 1 hour at room temperature. Blots were incubated with primary antibodies diluted in 3% BSA in TBST overnight at 4°C, washed with TBST three times for 10 minutes and blotted with secondary antibodies for 1 hour at room temperature.

After washing blots with TBST three times for 10 minutes, they were visualized with the LI-COR Odyssey® Infrared Imaging System and software at 169 µm (LI-COR Biosciences).

### Biotin-streptavidin affinity purification, mass spectrometry and data analysis

For the BioID experiments, HEK293T cells stably expressing V5-BirA* and V5-BirA*-ENKD1 were generated with lentiviral transduction. For large scale pulldowns, 5×15 cm plates were grown in complete medium supplemented with 50 μM biotin for 18 h. After biotin treatment, cells were lysed in lysis buffer (20 mM HEPES, pH 7.8, 5 mM K-acetate, 0.5 mM MgCI2, 0.5 mM DTT, protease inhibitors) and sonicated. An equal volume of 4°C 50 mM Tris (pH 7.4) was added to the extracts and insoluble material was pelleted. Soluble materials from whole cell lysates and centrosome enriched extracts were incubated with Streptavidin agarose beads (Thermo Scientific) overnight. Beads were collected and washed twice in wash buffer 1 (2% SDS in dH2O), once with wash buffer 2 (0.2% deoxycholate, 1% Triton X-100, 500 mM NaCI, 1 mM EDTA, and 50 mM HEPES, pH 7.5), once with wash buffer 3 (250 mM LiCI, 0.5% NP-40, 0.5% deoxycholate, 1% Triton X-100, 500 mM NaCI, 1 mM EDTA and 10 mM Tris, pH 8.1) and twice with wash buffer 4 (50 mM Tris, pH 7.4, and 50 mM NaCI). 10% of the sample was reserved for western blot analysis and 90% of the sample to be analyzed by mass spectrometry was washed twice in 50 mM NH_4_HCO_3_.

Mass spectrometry analysis was done as described (Gurkaslar et al., 2020). Raw data of three biological replicates for V5-BirA*-AURKA and three biological replicates for V5-BirA* were analyzed to identify proximity interactors of ENKD1. For control datasets, previously published datasets from V5-BirA*-expressing cells were used (Arslanhan et al., 2021). The data were analyzed with SAINTExpress (Teo et al., 2014). High confidence interactors were defined as the ones with BFDR score smaller than 0.03. GO terms were determined using g-Profiler. Interaction maps were drawn using Cytoscape.

### *In vitro* microtubule sedimentation assay

For sedimentation assays performed with whole cell lysates, HEK293T cells transfected with GFP-ENKD1 were lysed in in BRB80 buffer (80 mM PIPES pH 6.8, 1 mM EGTA, 1m M MgCl_2_) supplemented with protease inhibitors (10 µg/ml LPC, 1 mM PMSF and 10 µg/ml aprotinin) at 4°C for 30 minutes and centrifuged at 13.000 R.P.M. for 15 minutes. Lysate was further cleared by centrifugation at 90.000 R.P.M. for 5 minutes at 4°C with TLA100 rotor (Beckman). 100 µg cell lysate was brought to a final volume of 250 µl BRB80 supplemented with 1 mM GTP. Lysate was incubated with 25 µM taxol stepwise at 30°C for 30 minutes. Lysate was centrifuged at 55.000 R.P.M. for 10 minutes at 30°C in a TLA100 rotor (Beckman) through a 125 ul 40% glycerol cushion prepared from BRB90 buffer supplemented with 1 mM GTP. Pellets were resuspended in the same final volume as the input and equivalent volumes of pellet and supernatant were separated by SDS-PAGE and processed by immunoblotting.

For sedimentations assays performed with purified proteins, MBP-ENKD1 and MBP were expressed in Rosetta BL21 cells at 30C for 18 h. Bacteria expressing MBP-fusion proteins was lysed by sonication with 1mg/ml lysozyme in lysis buffer (20mM Tris pH:7.6, 250mM NaCl, 100mM KCl, 1mM DTT, 1 mM EDTA, 5% glycerol, 10 µg/ml LPC, 1 mM PMSF) and proteins were purified by amylose beads (New England Biolabs) and eluted in wash buffer (20mM Tris pH:7.6, 250mM NaCl, 100mM KCl, 1mM DTT, 1mM EDTA) with 10mM maltose. Purified bovine brain tubulin and purified proteins were precleared at 90.000 R.P.M. for 5 minutes at 4°C with TLA100 rotor. Cleared tubulin was polymerized by first incubating with 20 µM taxol stepwise at 37°C for 30 minutes followed by incubation at room temperature overnight. 0.75 nmol of purified protein was brought to final taxol concentration of 20 µM, mixed 1:1 with polymerized tubulin and incubated at 30°C for 30 minutes. Samples were loaded onto 40% glycerol BRB80 cushions and centrifuged at 55.000 rpm for 10 min with TLS55 rotor. Equivalent volumes of supernatant and pellet fractions were resolved by SDS-PAGE.

### In vitro microtubule bundling assay

Fluorescent MTs were polymerized at 2 mg/ml by incubating tubulin and rhodamine labeled tubulin (Cytoskeleton Inc.) at 10:1 ratio in BRB80 with 1 mM DTT and 1mM GTP for 5 min on ice, then preclearing by centrifuging for 10 min at 90000 rpm at 2°C. Cleared tubulin mixture was polymerized at 37°C by adding taxol and increasing concentration stepwise, with final concentration of 20 µM. MTs were pelleted over warm 40% glycerol BRB80 cushion in TLA100 rotor at 70000 rpm for 20 min. After washes with 0.5% Triton-X100, pellet was resuspended in 80% of the starting volume of warm BRB80 buffer with 1mM DTT and 20 µM Taxol.

Bundling assays were performed as previously described (Tao et al., 2016). In short, 100 nM His-MBP-ENKD1 or MBP protein was mixed with 2 µM MTs and 20 µM taxol in buffer T (20mM Tris, pH 8.0, 150mM KCl, 2mM MgCl2, 1mM DTT, protease inhibitors) for 30 min rocking at room temperature. The reaction mixtures were transferred into a flow chamber under a HCl-treated coverslip, and unstuck proteins were washed out with the excess of buffer T. Bundling of fluorescent MTs was observed with a Leica SP8 confocal microscope and 63x 1.4 NA oil objective (Leica Mycrosystems). Experiment was repeated 3 times.

### *In vitro* microtubule turbidity assay

Turbidity experiments were performed with 15 µM Tubulin (Cytoskeleton, Inc.) and desired concentrations of proteins, 100 nM MBP, 100 nM His-MBP-TOGARAM1TOG3-TOG4 and increasing concentration of His-MBP-ENKD1. Purified proteins were incubated at 4°C in BRB80 buffer supplemented with 1mM DTT, 1mM GTP and 25% glycerol. Tubulin was thawed on ice, precleared in a TLA100 rotor at 90,000 rpm for 10 min at 2°C and immediately mixed with proteins in the 60μl total volume assembly reactions. Tubulin polymerization was monitored using Epoch2 microplate spectrophotometer (BioTek) at 37°C. The absorbance at 350 nm was recorded at 5-s intervals for 35 minutes.

For microscopy-based analysis of microtubule polymerization, turbidity experiment was performed as described above but with rhodamine labeled tubulin (Cytoskeleton, Inc.) composing 1/10 of final tubulin concentration. After 35 minutes, samples were removed from plate and MTs fixed by diluting 11-fold into BRB80 buffer with 1mM GTP and 25% glycerol plus 1% glutaraldehyde at 37°C for 3 min. Fixation was quenched by addition of Tris (pH 7.0) to a final concentration of 100 mM and subsequently diluted ∼4-fold into BRB80 buffer with 1mM GTP without glycerol.

Microtubules were sedimented onto poly-L-lysine coated coverslips by centrifuging through 20% glycerol BRB80 cushion at 4700 rpm for 2 h in S6096 rotor (Beckman). Coverslips were subsequently fixed with 20°C methanol, mounted with Mowiol, and imaged using Leica SP8 confocal microscope and 63x 1.4 NA oil objective (Leica Microsystems).

### Statistical tests

No statistical method was used to estimate sample size. Statistical results, average and standard deviation values were computed and plotted using Excel or Prism 7 (Graphpad) Comparison of two groups was done using a two-sided Student’s t-test.

The comparisons of more than two groups were performed using one- or two-way ANOVAs. n indicates the number of experimental replicates for each sample and condition. Two-tailed t-tests were applied to compare measurements. The graphs with error bars indicate Standard Deviation (SD) and the significance level were denoted as \**P* < 0.05, \*\**P* < 0.01, \*\*\**P* < 0.001, **** *P* < 0.0001.

## Data availability

All data generated or analysed during this study are included in the manuscript and supporting files.

## Acknowledgements

We acknowledge the Firat-Karalar lab members for insightful discussions regarding this work. This work was supported by European Research Council (ERC) grant 679140 ENF, European Molecular Biology Organization (EMBO) Installation grant 3622 and an EMBO Young Investigator Award to ENF, TUBITAK grant 119Z347 to ENF and Marie Sklodowska Curie Fellowship to JD.

## Author Contributions

Conceptualization, ENF, FT, JD.; Methodology, ENF, FT, JD.; Investigation FT, JD.; Resources, ENF, FT, JD.; Writing—Original Draft, ENF, FT.; Writing—Review & Editing, ENF, FT, JD.; Visualization, ENF, FT.; Supervision, ENF.; Project Administration, ENF.; Funding Acquisition, ENF, JD.

---

**Figure S1.**
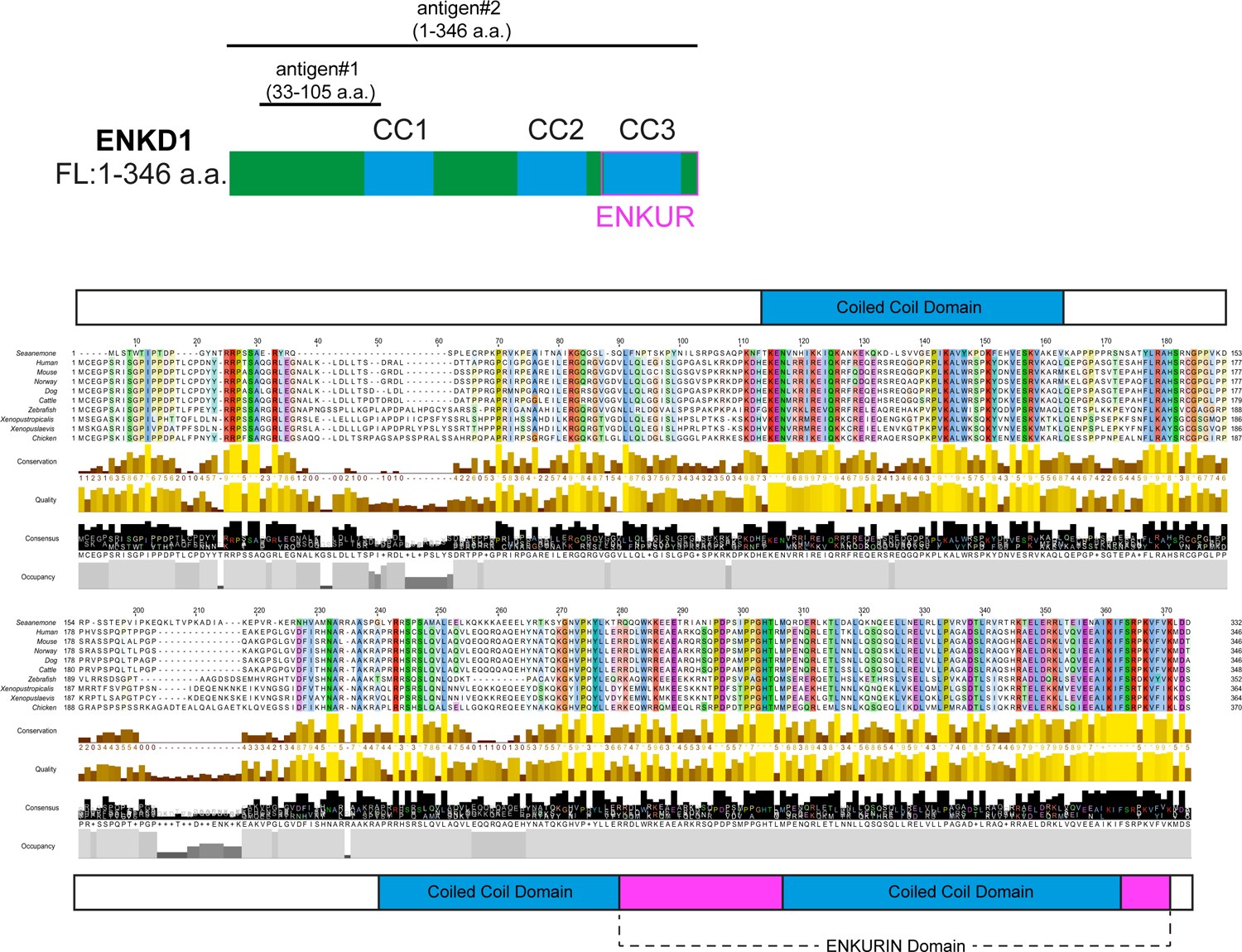
ENKD1 conservation analysis. Schematic representation of ENKD1 full length (FL, 346 a.a.). CC1, CC2, and CC3 indicate coiled coil domains and ‘ENKUR’ indicates the ENKUR domain. Human ENKD1 antibody (HPA041163, Sigma) recognizes antigen between the sequences of 33-105 a.a and rat polyclonal antibody was raised against purified MBP-fusion of full-length human ENKD1. Alignment of ENKD1 orhologous proteins from different organisms, colored by the Clustal X color scheme in Jalview. Conservation scores, locations of the predicted coiled-coils (CCs), enkurin domain were shown below the alignments.

**Figure S2.**
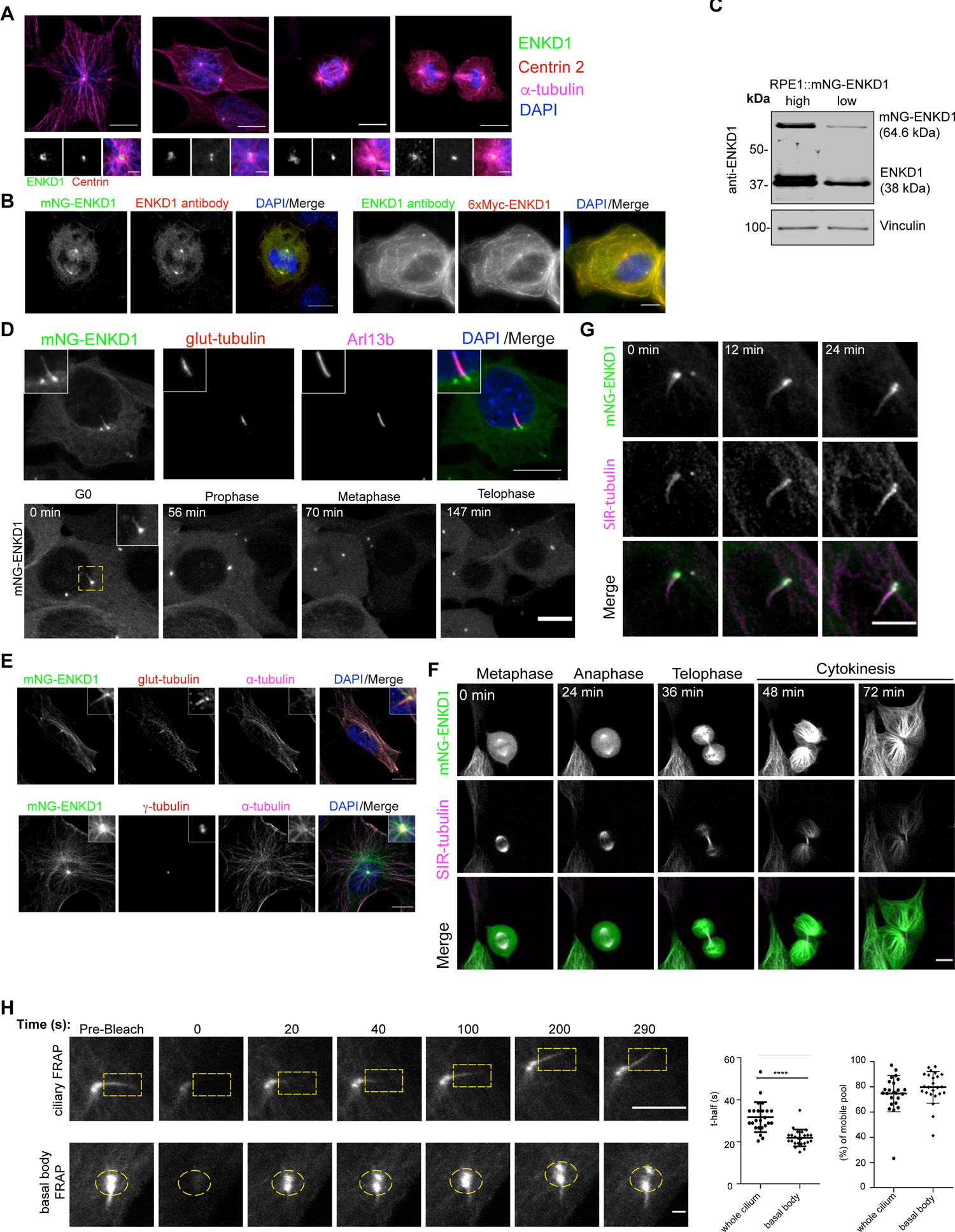
ENKD1 localization and dynamics in RPE1 and IMCD3 cells. (A) Endogenous ENKD1 localization in RPE1 cells. RPE1 cells were fixed and stained for rabbit ENKD1 antibody, centriole marker Centrin 2 and α-tubulin. Endogenous ENKD1 localizes to centrosome in interphase cells and spindle poles in dividing cells. (B) Immunoblotting of lysates prepared from low-expressing and high-expressing RPE1::mNG-ENKD1 cells with the ENKD1 antibody raised against 33-105 a.a. of ENKD1 and vinculin (loading control). (C) ENKD1 antibody recognizes mNG-ENKD1 in RPE1::mNG-ENKD1 stable cells and 6xMyc-ENKD1 in transiently transfected Hela cells. Cells were stained with ENKD1 antibody and antibody against the fusion tag. (D) Localization and dynamics of ENKD1 in low-expressing IMCD3::mNG-ENKD1 cells. (Top panel) Cells were serum staved for 24 h, fixed with PFA and stained for polyglutamylated-tubulin (glut-tubulin), Arl13b and and DAPI. Dynamic localization of ENKD1 throughout the cell cycle. (Bottom panel) Cells were stained with SiR-tubulin and analysed by time lapse confocal imaging. Still frames from Movie S3 were shown at the indicated time points to represent ENKD1 localization to the spindle poles and central spindle. Scale bar: 10 `μ`m (E) Localization of ENKD1 in high-expressing RPE1::mNG-ENKD1 cells. Cells were serum staved for 24 h, fixed with PFA and stained for alpha-tubulin (`α`-Tubulin), gamma-tubulin (`γ`-tubulin) and/or polyglutamylated-tubulin (glut-tubulin) and DAPI. Scale bar: 10 `μ`m. (F) Dynamic localization of ENKD1 throughout the cell cycle. High-expressing RPE1::mNG-ENKD1 were stained with SiR-tubulin and analysed by time lapse confocal imaging. Still frames from Movie S1 were shown at the indicated time points to represent ENKD1 localization to the spindle poles, spindle microtubules and central spindle. Scale bar: 10 `μ`m (G) Constitutive localization of ENKD1 at the primary cilium. IMCD3::mNG-ENKD1 cells were serum staved for 24 h, stained with SiR-tubulin and imaged for 24 min under serum starvation conditions. Still frames from Movie S2 were shown at the indicated time points to represent localization of ENKD1 to the primary cilium during the time course. Scale bar: 5 `μ`m (H) FRAP analysis of mNG-ENKD1 at the basal body and primary cilium. RPE1::mG-ENKD1 cells were serum starved for 24 h. Whole cilium and basal body indicated by yellow dashed rectangles and circles, respectively, was photobleached and cells were imaged for 5 min after photobleaching. Still images represent ciliary mNG-ENKD1 signal at indicated time points. Scale bar: 5 `μ`m for ciliary FRAP and Scale bar: 1 `μ`m for basal body FRAP. Half-time and mobile pool were calculated using recovery data. Half-time analysis of whole cilum is 31.74 (± 1.461, N=24); of basal body is 21.86 (±0.8181, N=24). Mobile pool percentage of whole cilum is 74.68 (± 2.945, N=24); of basal body is 79.68 (±2.559, N=24)

**Figure S3.**
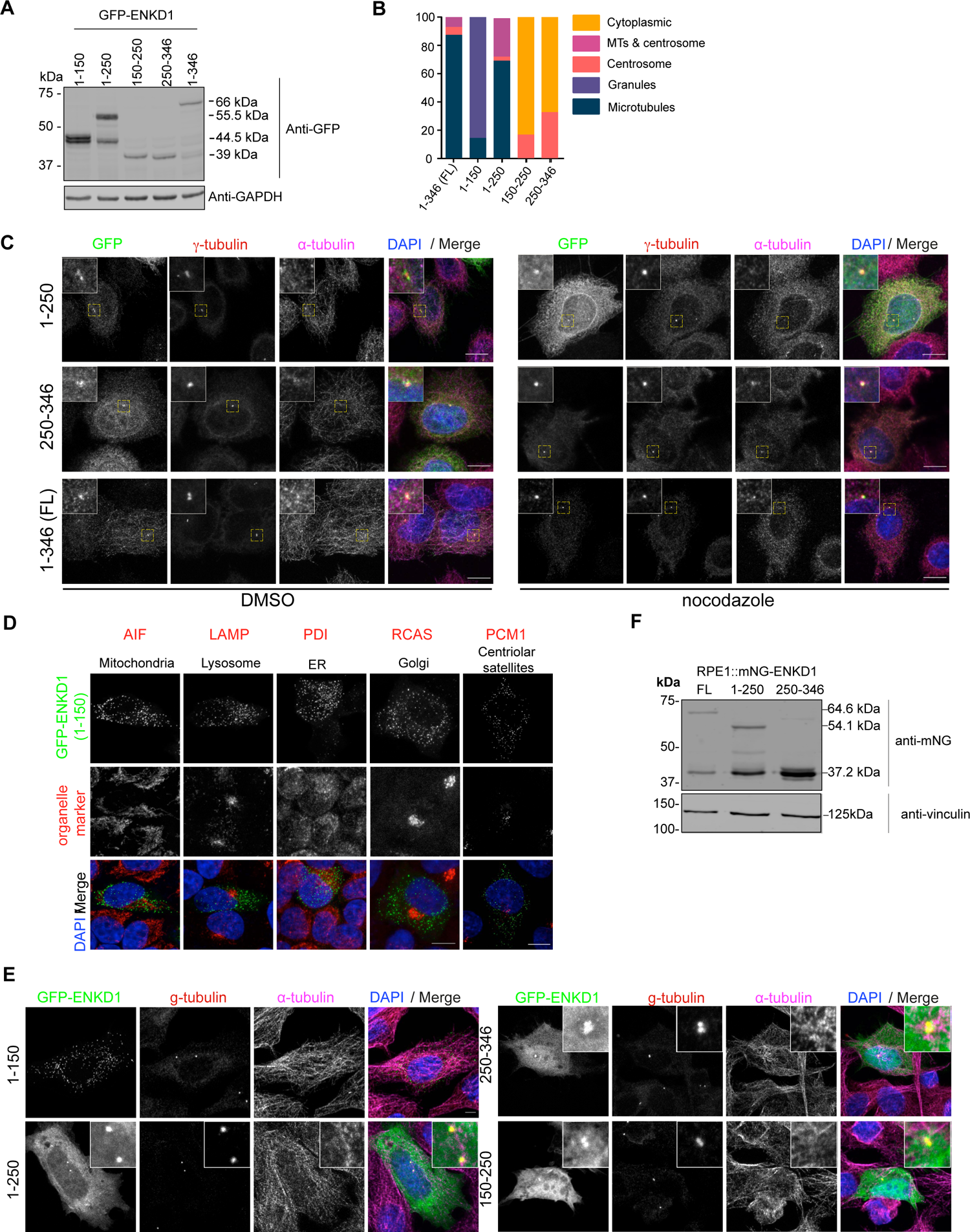
Localization analysis of ENKD1 deletion mutants. (A) Validation of expression of ENKD1 and its deletion mutants by immunoblotting. Lysates from HEK293T cells transfected with the indicated constructs were separated by SDS-PaGE and immunoblotted with antibodies against ENKD1 and GAPDH. Following are the GFP-fusions of the indicated proteins: ENKD1 (1-150) (44.5 kDa), ENKD1 (1-250) (55.5 kDa), ENKD1 (150-250) (38 kDa), ENKD1 (250-346) (38 kDa), and ENKF1 (FL) (66kDa). (B) Quantification of cellular localization of GFP-fusions of ENKD1 and its deletion mutans in transiently transfected RPE1 cells. Cells were fixed and immunostained with anti GFP, `γ`-tubulin and `α`-Tubulin antibodies. All transfected cells independent of their expression level were quantified based on their localization profile and data was plotted. (C) ENKD1 is a bona fide centrosome proteins. ENKD1 FL (1-346), (1-250), and (250-346) localizes to the centrosome in transfected HeLa cells treated with DMSO (vehicle control) or nocodazole for 1hr. Cells were fixed and stained for GFP, `γ`-tubulin and α-tubulin. Scale bar: 10 `μ`m (D) Relative localization of GFP-ENKD1 (1-150) relative to different organelle markers. HeLa cells transiently transfected with GFP-ENKD1 (1-150) were stained for antibodies against canonical organelle markers, which are AIF (mitochondrial marker), LAMP (lysosomal marker), PDI (endoplasmic reticulum marker), RCAS (Golgi marker), and PCM1 (centriolar satellite marker). Scale bar: 10 `μ`m. (E) Localization of GFP-fusions of ENKD1 and its deletion mutants in IMCD3 cells. (F) Validation of the RPE1 stable lines by immunoblotting. Lysates prepared from RPE1 cells stably expressing mNG ENKD1 FL, (1-250), and (250-346) constructs were separated by SDS-PAGE and immunoblotting with antibodies against mNG and vinculin. Expected sizes for the FL and deletion mutants are: ENKD1 (1-250) (54.1 kDa), ENKD1 (250-346) (37.2 kDa), and ENKD1 FL (64.6kDa).

**Figure S4.**
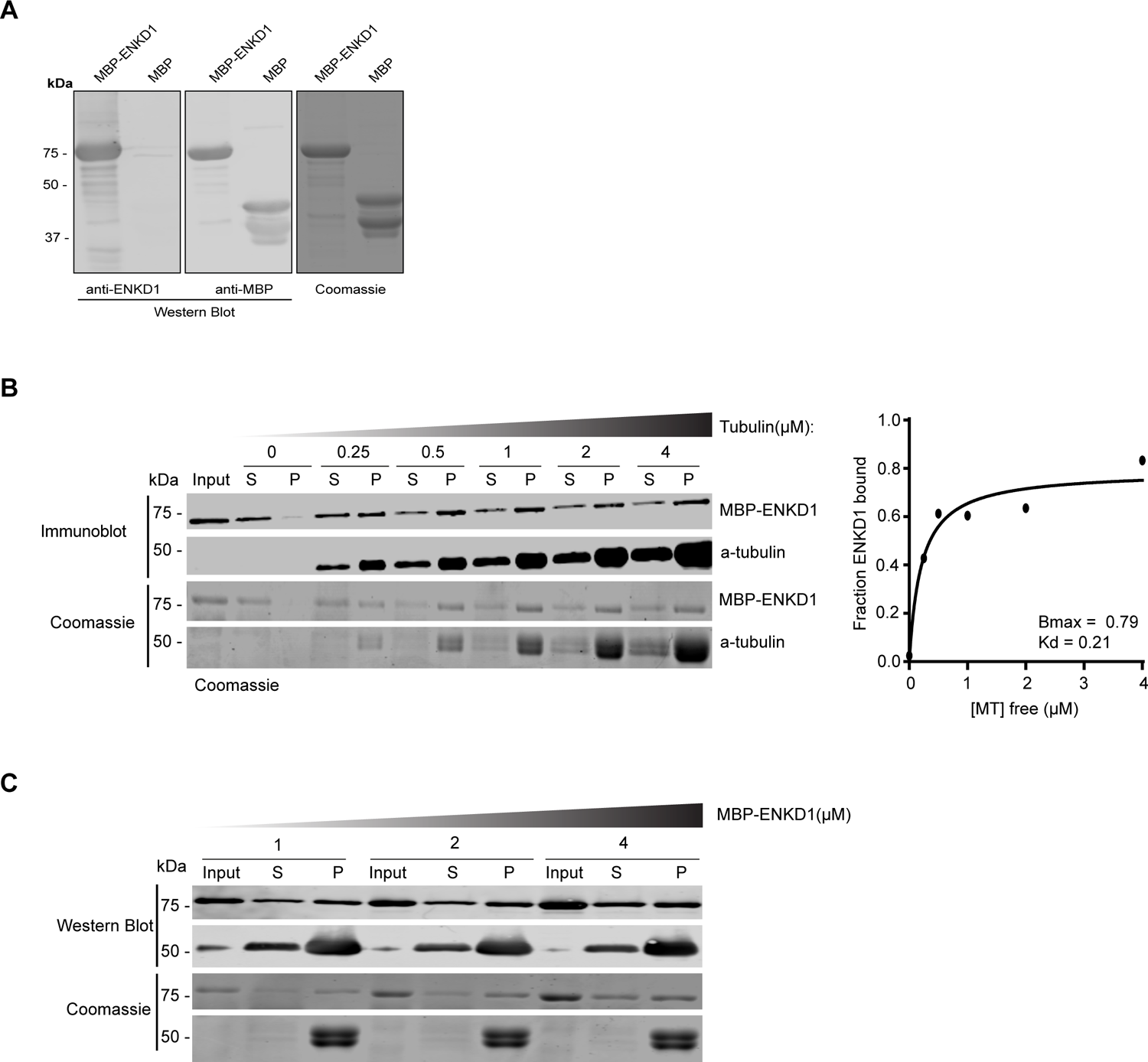
*In vitro* protein purification and microtubule co-sedimentations assays for MBP and MBP-ENKD1 (A) MBP and MBP-ENKD1 were expressed in *E. coli* and purified by amylose chromatography. Purified proteins were separated by SDS-PAGE gel and their purity and size were assessed by Coomassie Blue Staining and immunoblotting using antibodies against MBP or ENKD1. (B) *In vitro* microtubule co-sedimentation assay using 1 `μ`M of purified MBP-ENKD1 and 0, 0.25, 0.5, 1, 2 and 4 `μ`M Taxol-stabilized MTs. After incubation of proteins with MTs, reactions spun down (55,000 rpm for 10 min) on 40% glycerol cushion to pellet the polymerized MTs. Input, supernatant (S) and pellet (P) fractions were separated by SDS PAGE gel and proteins were detected by Coomassie Blue staining and immunoblotting using antibodies against ENKD1, and alpha-tubulin. The dissociation constant (*K*d) was determined from the best-fit curve and calculated by using Prism 6 software (one site-specific binding). Bmax=0.79 (±0.04); Kd=0.21 (±0.05). Data was derived from one experiment. (C) Microtubule co-sedimentation assay using increasing concentrations (1, 2, and 4 `μ`M) of MBP-ENKD1 and 1 `μ`M Taxol-stabilized MTs. Samples were prepared and analyzed as explained S5B.

**Figure S5.**
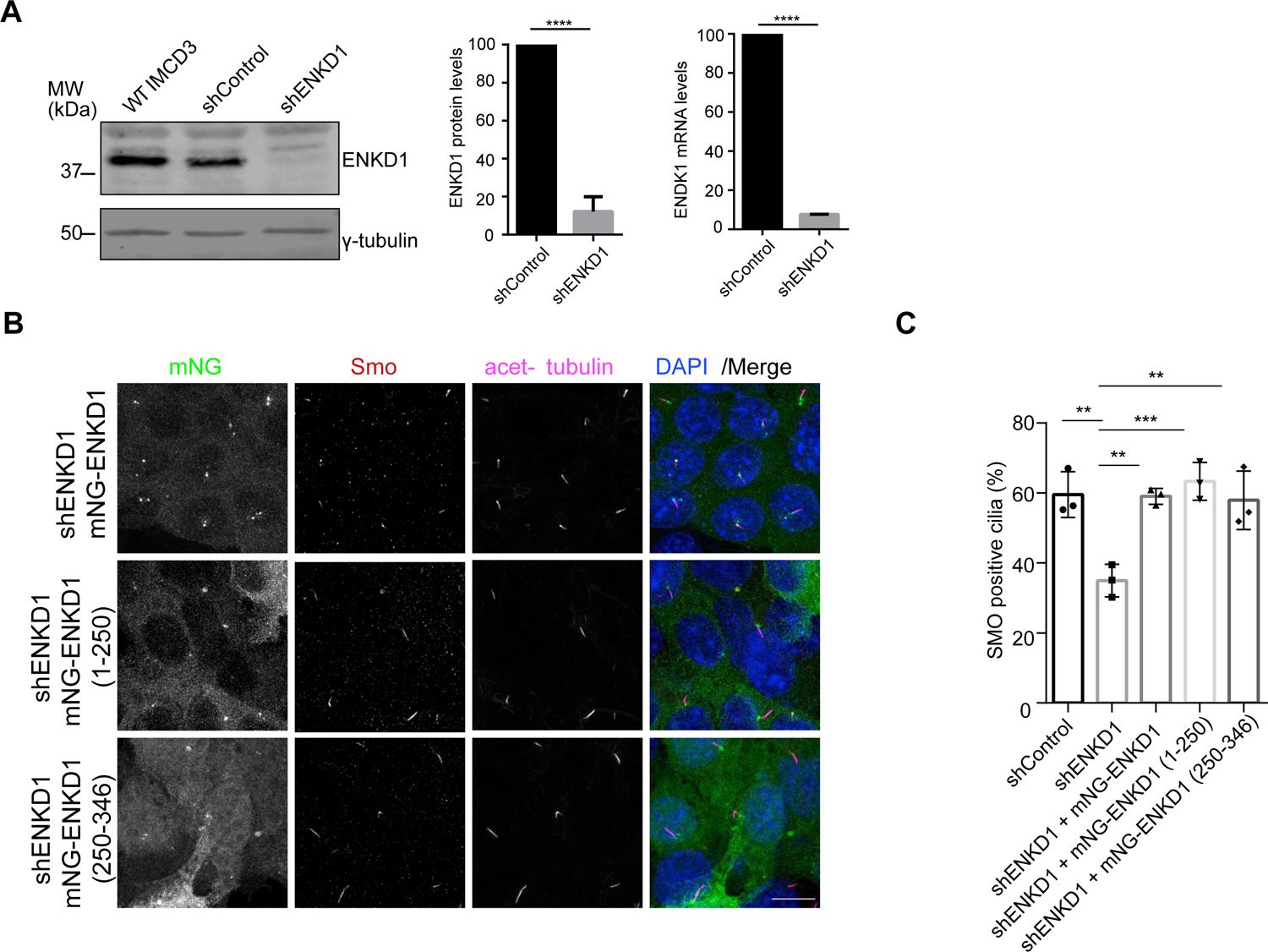
Validation of efficient ENKD1 depletion and the specificity of the Hedgehog pathway activation defects. (A) Validation of efficiency ENKD1 depletion using specific shRNAs. Immunoblot analysis of lysates prepared from wild type, control shRNA, and ENKD1 shRNA stably expressing IMCD3 cells. (First graph) Proteins were detected using antibodies against ENKD1 and `γ`-tubulin. Band intensities were measured using ImageJ. Mean shControl band intensity 100; mean shENKD1 band intensity 12.18 (±4.456). N=3. (Second graph) qRT-PCR showing reduced mRNA levels of ENKD1 in the IMCD3 cells stably expressing shENKD1. Normalized ENKD1 mRNA level in shControl is 100; in shENKD1 is 7.642 (± 0.06) N=2 (B) Rescue experiments for Hedgehog pathway activation defects. Stable expression of mNG-ENKD1, mNG-ENKD1 (1-250), and mNG-ENKD1 (250-346) in shENKD1 IMCD3 cells rescued ciliary SMO localization defect. Cells were serum-starved for 24 h, treated with 200 nM SAG for 4h, fixed and stained for SMO, acetylated tubulin (Ac. tub), and DAPI. (C) Quantification of S6B. % SMO localized to cilia in shENKD1 mNG-ENKD1 cells is 59.01; in shENKD1 mNG-ENKD1 1-250 cells is 63.31; in shENKD1 mNG-ENKD1 250-346 cells is 57.90 (N=3) Scale bar: 10 `μ`m.

## Movies

**1.** RPE1::mNG-ENKD1 cells were imaged every 2 minutes for 108 minutes. Scale bar:10 μm

**2.**100% confluent serum starved RPE1::mNG-ENKD1 cells were imaged every 2 minutes for 22 minutes. Scale bar: 1 μm.

**S1.** RPE1::mNG-ENKD1 cells were imaged every 3 minutes for 114 minutes. Scale bar:10 μm

**S2.** 100% confluent serum starved IMCD3::mNG-ENKD1 cells were imaged every 2 minutes for 24 minutes.. Scale bar: 5 μm

**S3.** IMCD3::mNG-ENKD1 cells were imaged every 7 minutes for 6h40 minutes. Scale bar:10 `μ`m

